# Pupil dilation as a marker of attention/effort in aging and mild cognitive impairment

**DOI:** 10.1101/2025.07.18.663475

**Authors:** Alina Zhunussova, Clare Loane, Elif Kurt, Grazia Daniela Femminella, Sabrina Lenzoni, Millie Duckett, Martina F Callaghan, Nikolaus Weiskopf, Raymond J Dolan, Robert Howard, Emrah Düzel, Dorothea Hämmerer

## Abstract

Pupil dilation (PD) can be easily measured and reflects responses to subjectively salient or cognitively demanding events. It therefore holds promise as a cognitive marker especially for individuals with mild cognitive impairment (MCI) or other neurodegenerative conditions with restricted abilities to respond in cognitive assessments. We assessed PD during two tasks, an oddball task for investigating attentional allocation and a Simon task which additionally allows for investigating cognitive effort in younger adults (YAs), older adults (OAs), and patients with MCI. PD is a useful marker for investigating attention and cognitive effort in MCI, as suggested by elevated PD to salient stimuli in particular of individuals with better attentional control in MCI patients, as well as YAs and OAs. Measurement of PD may serve as an easy-to-administer measure to assess changes in cognitive function in healthy aging and MCI.

## 1. BACKGROUND

Changes in pupil diameter, which are often unconscious, can reflect cognitive functions such as attention, effort or arousal [1,2]. More specifically, pupil size increases to salient stimuli which capture attention [3, 4, 5] or task-relevant stimuli which are to be met with a stronger a priori or top-down attentional focus [6,7]. A common experimental paradigm for inducing attentional pupil effects is the oddball task, where pupil size changes as participants respond to targets while ignoring irrelevant stimuli [8]. Beyond attentional effects, pupil dilation (PD) can also serve as a measure of cognitive load or (motor) effort [6,2]. For instance, pupil size has been shown to increase during more effortful compared to less effortful conditions in a digit-span working memory task [9,10], or as an indicator of motor effort and internal state changes during motor tasks [11,12].

While the full picture of the underlying mechanisms of cognition-related PD in humans is still unclear, recent studies point to subcortical neuromodulatory systems as a potential contributor. In particular, the rapid phasic PD, typical for pupil effects during cognitive tasks, has been associated with phasic activations in the noradrenergic locus coeruleus (LC) [13,14]. Indeed, stimulating the LC in monkeys has been shown to elicit a phasic PD [13,15,16], presenting strong evidence for a link between LC firing and PD. Based on LC imaging studies in humans [17,8,18] and animals [19,20,21], it is reasonable to assume that this is also the case for attentional and effort-related events. However, stimulation of other brain areas also results in phasic PD, suggesting that the link between LC firing and PD is not exclusive [13]. Indeed, a recent functional magnetic resonance imaging (MRI) study in humans showed that correlates of PD can be found in several brainstem centres [22,23]. Without intervention studies in animals which isolate specific PD-controlling (brainstem) structures, it is therefore currently unclear which brain structures are indispensable for driving PD and which might only contribute. Moreover, being able to establish these mechanisms in animal models might not translate completely to the processes underlying human PD. Given the small LC size and brainstem location, only a limited number of human neuroimaging studies have examined the relationship between LC activity and pupil to test animal-derived theories [22,24,23,8]. Specifically, Murphy [8] showed that the LC showed enhanced neural activity for rare target stimuli compared to standard stimuli in an oddball paradigm, thus supporting the contribution of the LC to attention-related PD in humans. Future pharmacological imaging studies might contribute to further disentangling brain structures indispensable for controlling PD in humans.

Taken together, the neural substrates of PD in animal models and humans are not fully understood. Nonetheless, given the consistent evidence for selective PD during distinct cognitive processes, such as the detection of salient events, the allocation of attentional resources, or effortful processing, pupillometry can be used to provide valuable insights into human cognitive processes. Moreover, despite providing only one data stream, PD can differentiate early versus late responses to specific cognitive events and distinguish multiple processes (e.g., emotional salience and memory recognition) [25,26]. As such, the ability to assess PD during cognitive tasks provides a fairly informative and comparatively easy-to-use method with minimal burden, even for cognitively impaired participants. Pupillary responses can be recorded at rest as well as during cognitive tasks and require no conscious effort of responding, making them suitable for unobtrusive cognitive assessments in e.g., in older adults (OAs) with mild cognitive impairment (MCI) or patients with Alzheimer’s disease (AD). Substantial deficits in executive control and attention characterize MCI and AD symptomatology [27,28,29], impacting daily functioning. Here, we explore the potential of using PD to characterize attentional functions in MCI during two executive attentional control tasks across MCI, age-matched controls, and younger adults (YAs). In doing so, we hope to add to studies that have used PD to characterize MCI- and AD-related changes in working memory [30,31], semantic processing [32], and recognition memory [33].

## 2. METHODS

### 2.1 Aims and hypotheses

The aim of this study is to investigate the sensitivity of pupillary responses to reveal differences in top-down attentional modulation and/ or cognitive effort in MCI patients compared to healthy age-matched adults. This will be tested using two tasks, an auditory and visual oddball task which allows for investigating attentional allocation to infrequent oddball stimuli [8], as well as a Simon task which additionally allows for investigating cognitive effort required to exert inhibitory control when navigating incongruent response inputs [34]. Additionally, to understand whether interindividual differences in LC degeneration are associated with PD or cognitive function in aging and MCI, as suggested by recent studies [35,36,37], we assess the structural integrity of the LC using neuromelanin-sensitive MRI in participants undergoing pupillometric recordings. While our main focus is on comparing PD in age-matched healthy OAs and MCI patients, we have also included a somewhat smaller sample of healthy YAs to serve as a reference for age-related differences in PD.

We make the following predictions:

1. OAs and patients with MCI show a similar pattern of pupillometric effects across task conditions, suggesting a similar degree of engagement of PD in cognitive processing during two attentional and cognitive control tasks.
2. For the oddball task assessing attentional allocation, we expect an increase in PD for the oddball compared to the standard stimuli [38] across both (auditory and visual) modalities, reflecting top-down attentional modulation to infrequent stimuli. Given the known decline in attentional executive control in MCI [39], we expect that this effect will be greater in OAs compared to MCI. Similarly, we hypothesize that OAs will demonstrate better behavioral performance than MCI patients, with higher accuracy rates and faster reaction times (RTs).
3. For the Simon task assessing cognitive control, we expect increased pupil responses to incongruent as compared to congruent stimuli reflecting greater top-down cognitive effort required to resolve stimulus-response conflicts. Given the age-related decline in executive functions [40,41], we hypothesize that the ability to engage attention and mobilize cognitive resources as evident in PD should be comparatively reduced in MCI compared to OAs. Moreover, both OAs and patients with MCI are expected to show the classic Simon effect, with longer RTs and lower accuracy for incongruent compared to congruent stimuli, with this effect being more pronounced in MCI patients as compared to OAs.
4. Within groups and tasks, we expect that greater PD, reflecting greater attentional focus or cognitive control, will be positively related to behavioral performance, i.e., shorter RTs and higher accuracy rates across individuals. In addition, higher LC integrity is expected to correlate with better cognitive performance and greater PD in OAs and MCI patients.

### 2.2 Participants

The data presented here is part of a larger study including behavioral, PD, functional MRI and structural MRI assessments. A total of 93 participants took part in the larger study, including 30 YAs, 30 healthy OAs, and 33 patients with MCI (see Table S1 in the Supplement). As the main focus of the study was to compare OAs and age-matched MCI participants, YAs were not routinely included in the PD recordings, especially for the oddball task. Therefore, whenever the number of participants in YAs was too small for statistical assessment, we report the results only descriptively to allow for an assessment of the typicality of the effects in comparison to existing pupillometry results in YAs. YAs were recruited via email from a participant panel at the UCL Institute of Cognitive Neuroscience. OAs were recruited via advertising and clinical teams when accompanying relatives or friends with MCI/ AD to a memory clinic. MCI patients were recruited through the Join Dementia Research participant database and local memory clinics at Camden and Islington Healthcare Trust and North East London Healthcare Trust. All participants were right-handed, had normal or corrected-to-normal vision, and had no history of psychiatric disorders. MCI in patients was confirmed through neuropsychological evaluation, that is a score at least 22 or above on the Mini Mental State Examination (MMSE) [42] as well as a diagnosis by a trained clinician. Healthy OAs and MCI patients were age matched and had to be within the age range from 60 to 85 years. YAs were between 20 to 30 years old. Caregiver-rated assessments were obtained for 27 of the 33 MCI patients, as 6 were not accompanied by caregivers. The mean age of caregivers was 67.81 years (SD = .42; age range = 35-84), with 6 males and 21 females.

As detailed below, data from several participants had to be excluded from the analysis due to four main reasons: (1) technical issues with the eye tracker, resulting in missing or unusable data, (2) difficulties in completing the tasks, especially in MCI patients, and (3) insufficient PD data due to excessive artifacts (e.g., blinking) in the case of the oddball task and (4) challenges of PD acquisition in the MR environment in the case of the Simon task related to insufficient signal strengths in infrared recordings.

Specifically, for the auditory and visual oddball tasks, we had to exclude: 16 OAs (12 due to technical issues with the eye tracker set-up in the eye tracking lab (hardware malfunctions, specifically issues with connection cables in the eye tracking lab), resulting in missing or unusable data, 2 due to difficulty in completing the task, resulting in no behavioral data being recorded, and 2 due an insufficient number of PD trials (less than three in one condition) due to excessive artifacts (e.g., blinking) in the PD data in one or both of the tasks); 10 patients with MCI (5 due to technical issues with the eye tracker, 4 due an insufficient number of PD trials, and 1 due to difficulty in completing the task; 5 YAs were excluded from statistical analyses, 2 because of insufficient numbers of PD trials, 2 due to technical issues with the eye tracker, and 1 due to difficulty in completing the task. Given the limited recruitment of YAs, this resulted in a small subsample (n=7) of YAs for the oddball tasks and their data being presented descriptively for reference only. The final sample for both the visual and auditory oddball tasks included 14 healthy OAs and 23 individuals with MCI.

PD during the Simon task was acquired concurrent to fMRI assessments (not reported here). PD assessments were therefore acquired in the whole sample of 93 participants. However, PD is more challenging to acquire in the MRI due to several factors. Eye tracking quality is often compromised due to a) shadows from the head coil which obscure pupil recordings depending on the position of the head in the coil, and b) a weaker pupil signal due to the position of the illuminator and camera behind the scanner bore compared to standard laboratory eye-tracking setups. In addition, head movements during recording can interfere with pupil signal acquisitions. For the Simon task, across the whole sample of 93 participants, a total of 31 participants were excluded: 8 YAs (6 due to challenges with acquiring PD in MRI setting, resulting in missing or unusable data, 1 due to difficulty in completing the task, and 1 due to unusually slow performance (RTs greater than three standard deviations from the group mean), 10 OAs (8 due to challenges with acquiring PD in the MRI setting and 2 due to difficulty in completing the task), and 13 patients with MCI (10 due to challenges in acquiring PD in the MRI setting and 3 due to difficulty in completing the task). The final sample for the Simon task consisted of 22 healthy YAs, 20 healthy OAs, and 20 individuals with MCI.

Although sample sizes are at the lower bound, low sample sizes are not uncommon in eye tracking studies [32,33,43] which exhibit typically reliable PD effects in well-established attentional and cognitive control tasks as employed here [8,44]. Moreover, to explore whether our study might have yielded spurious results given small and heterogeneous sample sizes, we explore the consistency of PD effects across individuals per group (Figures S1-S3 for the oddball task and Figures S4 and S5 for the Simon task in the Supplement). On the oddball tasks, larger PDs for oddball as compared to standard stimuli were observed in 92-100% of YAs, OAs or MCI patients; on the Simon task, PD was larger for the incongruent condition in 68% of YAs, 85% of OAs, and 90% of MCI patients. The high consistency of these well-established PD effects therefore suggests that our results are representative despite somewhat reduced sample sizes.

Finally, age matching was reconfirmed after exclusions for both the oddball task, t(29.89) = −1.61, p = .11), with mean ages of 69.92 and 73.91 for OAs and MCI patients, respectively, and the Simon task, t(37.63) = −.84, p = .40), with mean ages of 71.2 and 73.2 years for OAs and MCI patients, respectively.

### 2.3 Procedure

Participants and/ or caregivers completed neuropsychological and clinical questionnaires in a first session prior to MRI and PD assessments. All participants completed the National Adult Reading Test [45], Addenbrooke’s Cognitive Examination (ACE) [46], including the MMSE. For clinical assessments, YAs and OAs completed the Hospital Anxiety and Depression Scale [47], and the Pittsburgh Sleep Quality Index (PSQI) [48]. MCI patients and/ or caregivers completed the PSQI, Bristol Activities of Daily Living [49], Cornell Scale for Depression in Dementia [50], and the Neuropsychiatric Inventory [51]. The ACE is cognitive test designed to assess cognitive function in five domains, with higher scores indicating better cognitive function: attention (18), memory (26), verbal fluency (14), language (26), and visuospatial ability (16). A total score of 85 or less indicates cognitive impairment [52]. The revised version of the ACE (ACE-R) includes the MMSE, with a total score of 30. Scores below 24 may indicate cognitive impairment. In this paper, we present results for two domains of the ACE-R – attention/ orientation and memory - as well as the MMSE because of their diagnostic relevance to MCI, since these tests target cognitive domains known to be affected in the pathological progression of Alzheimer’s disease.

PD during the Simon task were recorded in the MR, together with an emotional memory task (not reported here). The oddball tasks were performed on the afternoon outside of the MR in an acoustically shielded room with controlled illumination. Each participant was reimbursed with £10 per hour for the time spent. Written informed consent was obtained from all participants and their carers prior to participation in the experiment. The study was conducted in accordance with the Declaration of Helsinki, ethical approval was granted by the local ethics committee (University College London ethics reference number 17/0091).

### 2.4 Eye-tracking recording

Pupillometric data were recorded using a laboratory-based (oddball task) or scanner-mounted (Simon task) infrared eye tracker (EyeLink 1000, SR Research). Participants’ right eye pupil size was tracked at 1000 Hz and illumination levels of 100%. The lighting in the test room remained constant throughout the tasks. The eye tracker was calibrated prior to each task using a 6-point or 9-point calibration.

### 2.5 Tasks

#### Oddball Task

The task was programmed using MATLAB R2015a (version 8.5.0.197613, The MathWorks, Inc, 2015) and Cogent 2000 software. It was presented on a 19-inch computer screen with a resolution of 1024×768 pixels. Participants were seated at a distance of 80 cm from the screen, with their head stabilized by a chin rest and head rest mounted on the desktop. All stimuli, including background and fixation crosses were luminance adjusted to 50% brightness to control for luminance-related effects on PD. For the auditory oddball task, the stimulus was presented binaurally through headphones while participants were fixating a fixation cross in front of a checkered background. The oddball was a high frequency tone (2000 Hz) and the standard stimulus a low frequency tone (1000 Hz), the loudness of both was adapted to the individual’s hearing ability prior to the task (Figure 1A). For the visual oddball task, the oddball stimulus was a triangle and the standard stimulus was a square, both filled with a red color and outlined with a dark border. Stimuli were presented in front of a gray checkered background (Figure 1B). Responses to the oddball task were given using a specific button on a 4-alternative button box. Recordings in the LC in animal studies suggest that the LC is also active during response initiation [53]. In order to maximize PD to oddball stimuli, participants were therefore asked to only respond to oddball stimuli (cf. 8 for a similar approach in a study recording LC responses during an oddball task). For auditory and visual oddball task, a trial began with a jittered fixation cross (250-750 ms, mean duration 500 ms), followed by a visual or auditory stimulus which lasted 2000 ms, followed by a response period of 2000 ms. Participants were instructed to let their index finger rest on the button after the practice to avoid pressing the wrong button.

**Figure 1.**
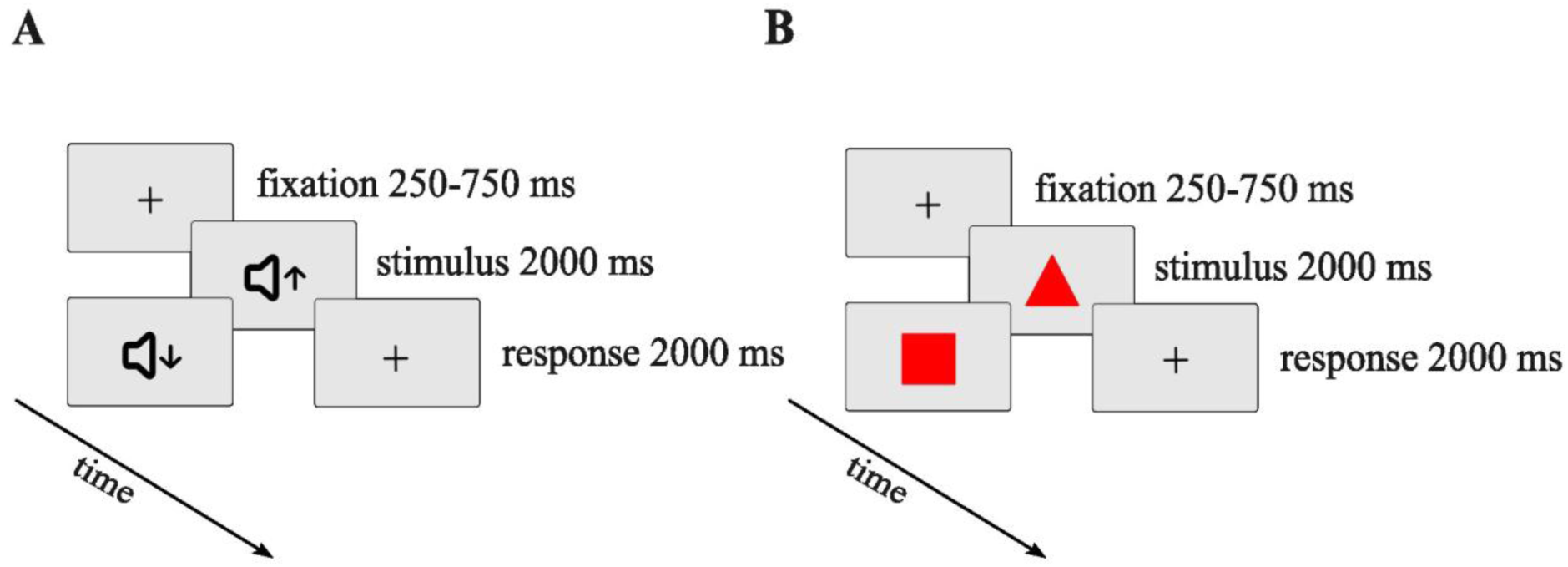
Graphical representation of the trial structures and timings for both types of oddball tasks. (A) Auditory Oddball Task: Participants were presented with either the oddball or the standard stimulus (high or low frequency tone, respectively). (B) Visual Oddball Task: Participants were presented with either the oddball or standard stimulus (triangle or square, respectively).

The oddball task consisted of 8 alternating blocks, beginning with the visual oddball task and then alternating between the task types. Participants were reminded of the task instructions at the beginning of each block. Each block consisted of 5 oddball trials and 20 standard trials, resulting in a total number of 20 oddball and 80 standard trials per task type. Participants were asked to respond to the oddball stimuli only. All participants completed a short practice session for both stimulus types.

#### Simon task

The task was performed in an MRI scanner and presented on a 19-inch computer screen with a resolution of 1024×768 pixels, viewed through a mirror attached to the head coil. Stimuli were solid red or green arrows presented in the center of the screen, preceded by a gray fixation cross. As for the oddball task, checkered background, fixation cross and stimuli were luminance controlled to 50% brightness. A trial began with a jittered fixation cross (250-750 ms, mean duration 500 ms) followed by an arrow stimulus for 2500 or 4500ms (based on average RTs during practice). A response was counted if it was executed during stimulus presentation. MR-compatible button boxes in each hand were used for response execution with left and right index fingers. YAs (n = 22) and OAs (n = 22) had an average response deadline of 2500 ms, whereas MCI patients (n = 20) had an average response deadline of 3000 ms, with five patients requiring an extended deadline of 4500 ms. Participants were asked to respond differently to the green and red arrows. For green arrows (congruent condition), participants pressed the button with the hand corresponding to the direction of the arrow (i.e., a right-pointing arrow required a right-hand response) (Figure 2A). For red arrows (incongruent condition), participants pressed the button with the hand opposite to the direction of the arrow (right-pointing arrow required a left-hand response) (Figure 2B). Instructions and a practice session of 12 trials were completed outside the MRI scanner on a desktop computer. The main test session consisted of 200 trials, of which 160 trials were congruent stimuli (green arrow) and 40 incongruent trials (red arrow). Both trial conditions consisted of an equal number of left pointing and right-pointing arrows.

**Figure 2.**
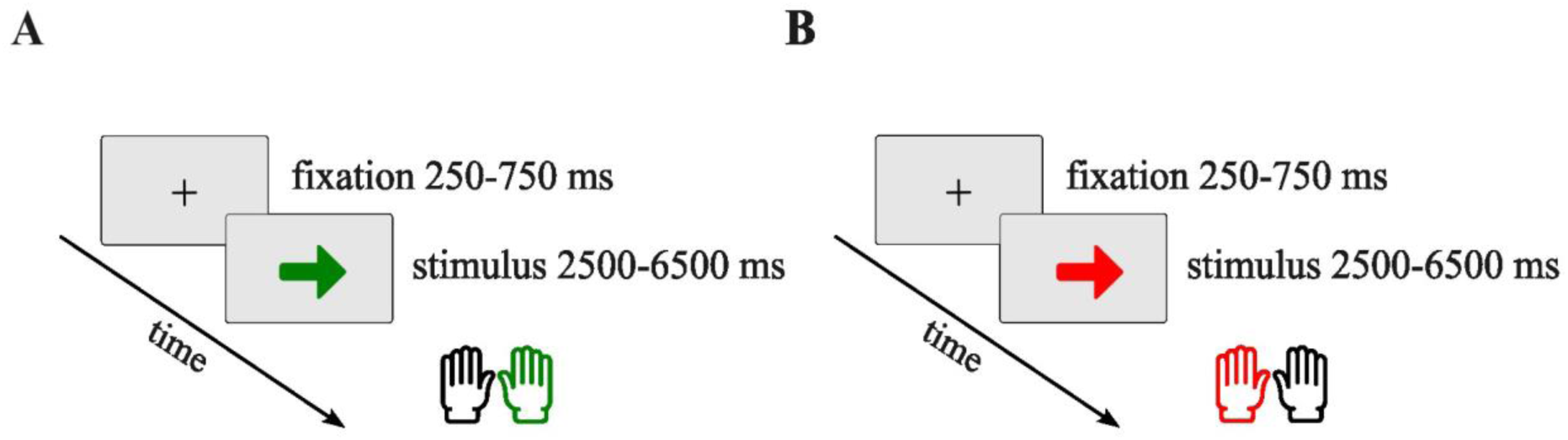
Graphical representation of the trial structures and timings in the Simon task. (A) Congruent trials. For the green arrow, participants had to respond with the hand that was congruent with the arrow direction. (B) Incongruent trials. For the red arrow, participants had to respond with the hand that was incongruent with the arrow direction.

### 2.6 Acquisition and pre-processing of pupillometric data

PD was recorded continuously throughout the experiment and segmented in a time window of 200 ms before and 2500 ms after relevant events (i.e., oddball or target onset, onset of congruent or incongruent stimulus) for assessing condition effects in phasic PD. To enable comparisons of phasic PD regardless of individual pupil size variations and distance to the eye tracker camera, segmented pupil data were concatenated and z-scored per individual before assessing conditions effects [54]. Phasic pupil responses were assessed relative to a 200ms baseline prior to stimulus presentations, that is, the mean PD in the baseline window was subtracted from the event-related PD for each trial. Moreover, given that intraindividual stability and noise in physiological processes can be expected to be altered in aging or MCI [55], an indicator of measurement noise in PD was calculated based on variations in PD across time for each trial in the 200 ms pretrial window, averaged across all trials per participant, yielding an individual-specific measure of PD variations in the absence of task-related events (referred to here as ‘baseline noise’) [54]. In case of group differences in baseline noise in PD recordings, estimates of baseline noise were included as covariate. Missing data due to artefacts and blinks were detected in MATLAB 2015b using the MATLAB-based toolbox FieldTrip (http://www.fieldtriptoolbox.org/). Missing data was replaced by linear interpolation using the interp1.m implemented in MATLAB. As missing data due to blinks or artefacts is typically preceded and followed by corrupted data (e.g., due to the eyelid closing and opening), data was interpolated within time periods of 200 ms or 30 ms before and after, large or small artifacts, respectively. Interpolated data were visually inspected and trials were excluded if fewer than three trials were available in either condition (see Table S2 in the Supplement for trial numbers and group comparisons).

To compare condition effects, the mean PD for each participant was estimated by averaging the PD for the auditory oddball and Simon tasks during the 1000–2000 ms time periods. Since the visual stimulus elicited a faster pupil response than the auditory stimulus during the oddball task, the mean PD for the visual task was computed within a time range of 500–1500 ms post stimulus onset.

### 2.7 Structural LC imaging

Scanning was performed with a 3T Siemens TIM Trio System and a 32-channel head coil. LC volumes were assessed using high resolution (.6*.6*3mm, 20 slices) magnetization transfer-weighted (MT-weighted) 3D Gradient Echo images (FLASH), (TR = 24.50 ms, TE = 3.35-11.75 ms, FA = 12), averaged across 3 repetitions. Quality assurance checks were performed to identify image artifacts, and sequences were repeated when recording artifacts were detected. Likely due to neuromelanin deposits in the LC, the LC is visible as a bright structure in so-called neuromelanin-sensitive scans [56,57]. As deposits are accumulating within LC cells, LC brightness is therefore interpreted as an indirect indicator of the structural integrity of the LC [58]. The LC mask was determined using the consensus between two independent raters’ voxel-wise segmentations in ITK-SNAP. To calculate a measure of LC integrity, neuromelanin signal intensity was taken as a ratio score of signal intensity in the LC mask to signal intensity in a reference region without neuromelanin located nearby in the pons. LC contrast ratio (CR) was then computed by dividing the difference between the intensity of the LC signal and the reference signal by the reference signal intensity [59]. The size and location of the reference region used was determined based on previous studies [60]. Two independent raters (DH and MD) manually segmented the LC using ITK-Snap as previously described [36]. LC integrity was calculated as the ratio score of signal intensity in the conjunction mask across both raters relative to the signal intensity in a reference region located in the pons not assumed to contain NM [35,36]. Sørensen–Dice coefficients for inter-rater consistency across the three groups were .70 (YAs), .65 (OAs) and .57 (MCI), respectively. Signal brightness in three brightest voxels was averaged per participant and used as a more robust measure of LC integrity [61].

### 2.8 Data Analyses

Data quality control involved several steps. First, participants were excluded if they had missing pupil data, insufficient behavioral data, or failed to perform the task according to task instructions. Specifically, RTs smaller than 200 ms were considered implausibly fast and excluded from the analysis. RTs greater or smaller than three standard deviations from the group mean were considered potential outliers and removed. Given the right-skewed distribution, RTs were log-transformed for normalization. Finally, each participant’s aggregated PD data was visually inspected to identify subjects with consistent measurement abnormalities or signal instability across multiple trials.

Missing data on neuropsychological tests and LC integrity measures were imputed using the expectation-maximization algorithm, using age as a predictor. In total, out of the 93 participants, four (all MCI patients) for ACE-R total, four (all MCI patients) for ACE-R memory, four (all MCI patients) for ACE-R attention/ orientation, four (all MCI patients) for ACE-R verbal fluency, four (all MCI patients) for ACE-R language, four (all MCI patients) for ACE-R visuospatial, two (two MCI patients) for MMSE, and three subjects (one YAs and two MCI) were imputed for LC integrity. For the ACE-R assessment, ACE-R total showed means of 94.43 (SD = 4.75) for YAs, 96.03 (SD = 3.37) for OAs, and 78.45 (SD = 11.70) for MCI patients. ACE-R memory subscale means were 24.27 (SD = 2.11) for YAs, 24.63 (SD = 2.00) for OAs, and 15.79 (SD = 6.44) for MCI patients. ACE-R the attention/ orientation subscale showed means of 17.87 (SD = .43) for YAs, 17.80 (SD = .48) for OAs, and 15.21 (SD = 2.02) for MCI patients. ACE-R the verbal fluency subscale means were 12.57 (SD = 1.43) for YAs, 12.93 (SD = 1.14) for OAs, and 9.34 (SD = 2.80) for MCI patients. ACE-R the language subscale means were 24.17 (SD = 2.15) for YAs, 25.53 (SD = .77) for OAs, and 24.03 (SD = 2.38) for MCI patients. ACE-R the visuospatial ability subscale means were 15.43 (SD = 1.13) for YAs, 15.20 (SD = 1.09) for OAs, and 14.07 (SD = 2.72) for MCI patients. Average MMSE scores were 29.53 (SD = 1.07) for YAs, 29.43 (SD = .72) for OAs, and 25.19 (SD = 2.79) for MCI patients. LC integrity measurements prior to imputation showed means of .13 (SD = .03) for YAs, .16 (SD = .04) for OAs, and .10 (SD = .03) for MCI patients. The imputation procedure did not significantly alter the distribution of these initial values (see Table S1 in the Supplement for detailed statistics with imputed values).

Statistical analyses were performed with R version 4.4.1 (2024-06-14) (R Core Team, 2024). The following R packages were used: ggplot2 (3.5.1) for data visualization, car (3.1.3) for Levene’s test, afex (1.4.1) for ANOVAs, emmeans (1.10.4) for estimated marginals means and pairwise comparisons among the means, effectsize (0.8.9) for calculating effect sizes for t-tests, cocor (1.1.4) for Williams’ t-test and Fisher’s z-test. Significance levels were set at α = .05. Multiple pairwise comparisons were conducted using estimated marginal means (emmeans package) and adjusted using multivariate t-distribution correction to control family-wise error rate. Effect sizes were calculated using partial eta squared (η^2^p). For one-way ANOVAs, eta squared (η^2^) was used. Cohen’s d (d) was calculated to assess effect size for independent t-tests.

### 2.9 Statistical analyses

To verify age matching between the OAs and MCI patients, we performed a two-sample t-test, which confirmed no significant age differences between the groups (see Methods section 2.2, description of participants).

For PD on the oddball tasks, we performed a repeated measures ANOVA with two within-subject factors: stimulus type (two levels: standard, oddball) and task (two levels: auditory, visual) and a between-subject factor: group (two levels: OAs, MCI). For PD within each task, we performed a repeated measures ANOVA with a within-subject factor: stimulus type (two levels: standard, oddball) and a between-subject factor: group (two levels: OAs, MCI). Baseline pupillary noise as defined by temporal variations within the baseline window differed significantly between groups, t(35) = −2.02, p = .05, d = .68, with OAs showing lower baseline noise (M = .14) compared to MCI patients (M = .16). Given these group differences, we mean-centred and included baseline noise as a covariate in subsequent analyses. Levene’s test indicated homogeneity of variances across groups for baseline noise (F(1, 35) = 2.76, p = .10), mean PD for standard stimuli in the visual oddball task (F(1, 35) = .03, p = .86), mean PD for oddball stimuli in the auditory oddball task (F(1, 35) = 1.69, p = .20), and mean PD for standard stimuli in the auditory oddball task (F(1, 35) = .13, p = .71). However, this assumption was violated for mean PD for oddball stimuli in the visual oddball task (F(1, 35) = 5.15, p = .02). For behavioral assessments, we used a repeated measures ANOVA with a within-subjects factor: task (two levels: auditory, visual) and a between-subjects factor: group (two levels: OAs, MCI). In addition, we used two-sample t-tests to examine differences between the OAs and MCI groups in performance within each oddball task. Levene’s test indicated homogeneity of variances across groups for hit rate in the visual oddball task (F(1, 35) = .95, p = .33), discrimination accuracy in the visual oddball task (F(1, 35) = 1.52, p = .22), RTs in the visual oddball task (F(1, 35) = .05, p = .82), and RTs in the auditory oddball task (F(1, 35) = 0, p = .99). However, this assumption was violated for the hit rate in the auditory oddball task (F(1, 35) = 10.46, p < .01) and the discrimination accuracy in the auditory oddball task (F(1, 35) = 7.03, p = .01). In this case, the Welch two-sample t-test was used.

For PD on the Simon task, we used a repeated measures ANOVA with a within-subjects factor: condition (two levels: congruent, incongruent) and a between-subjects factor: group (three levels: YAs, OAs, and MCI). As in the oddball tasks, analysis of baseline pupil noise revealed significant group differences (F(1, 59) = 14.74, p < .001, η^2^p = .33). Post hoc comparisons revealed that YAs had lower baseline noise (M = .16) compared to both OAs (M = .19) and MCI patients (M = .22), while OAs also had significantly lower baseline noise compared to MCI patients. Therefore, baseline noise per individual was mean-centred and included as a covariate in the analysis of the pupil data. Levene’s test indicated homogeneity of variances across groups for baseline noise (F(2, 59) = 2.61, p = .08), mean PD during congruent trials (F(2, 59) = .06, p = .93), and mean PD during the incongruent trials (F(2, 59) = .22, p = .80). For behavioral assessments, we performed a repeated measures ANOVA with a within-subjects factor: condition (two levels: congruent, incongruent) and a between-subjects factor: group (three levels: YAs, OAs, and MCI). Levene’s test indicated homogeneity of variances across groups for RTs during correctly performed incongruent trials (F(2, 59) = 2.94, p = .06). However, this assumption was violated for accuracy during correctly performed congruent trials (F(2, 59) = 6.45, p < .01) and correctly performed incongruent trials (F(2, 59) = 4.54, p = .01), and for RTs during correctly performed congruent trials (F(2, 59) = 4.22, p = .01) and incorrectly performed incongruent trials (F(2, 37) = 7.40, p < .01).

We examined relationships between variables using Spearman’s correlations, analyzing associations between: (1) PD and behavioral performance, (2) PD and LC integrity, (3) task measures and cognitive screening measures. First, we conducted correlational analyses between the variables of interest. For those correlations that showed significant relationships, we performed outlier analyses using Cook’s distance to identify influential data points (outliers). Observations were considered highly influential and eliminated from the correlational analyses if their Cook’s distance exceeded 4/n (where n is the sample size of each group). For the single oddball task correlational analysis, an average of one case of YAs, one case of OAs, and two cases of MCI patients were excluded after being identified as highly influential outliers. For the Simon task correlational analyses, an average of .66 YAs, .33 OAs, and .66 MCI patients were excluded from each group across the six correlational tests. For the single correlational analysis of incorrectly performed incongruent trials in the Simon task, two participants were excluded: one from the OAs and one from the MCI patients. While Williams’ t-test was used to assess significantly different correlations within the same group, Fisher’s z-test was used to compare the difference between two correlations in two independent groups.

To identify relationships across individuals in the context of small sample sizes, we performed a multiple linear regression analysis with an interaction term to examine the relationship between PD for congruent/ incongruent trials and oddball/ standard stimuli and LC integrity as well as RTs for congruent/ incongruent trials and oddball/ standard stimuli and LC integrity. Our analysis focused only on older groups (OAs and MCI patients), as LC integrity in younger age groups is likely more dominated by interindividual differences in neuromelanin accumulation rather than cell loss [62]. A binary group variable was created, where 0 represented OAs and 1 represented MCI patients. The linear model was constructed with mean PD as the dependent variable and group and LC integrity as the independent variables. Additionally, the linear models for the RTs were constructed with RTs as the dependent variable and group and LC integrity as the independent variables. The analysis was performed using the lm() function in R.

## 3. RESULTS

### 3.1 Oddball task

#### 3.1.1 Behavioral results

##### Between tasks (visual oddball vs auditory oddball)

A repeated measures ANOVA (group × task) for accuracy revealed a group effect, with OAs showing higher accuracy than patients with MCI regardless of the stimulus modality, with a higher hit rate (OAs: M = .94; MCI: M = .87) (F(1, 35) = 8.38, p < .01, η^2^p = .19) and discrimination accuracy (hit rate minus false alarm rate) (OAs: M = .94; MCI: M = .85) (F(1, 35) = 7.46, p = .01, η^2^p = .17). In addition, our analysis revealed a main effect of task, indicating that participants performed better on the visual compared to the auditory oddball task, in terms of faster RTs (visual: M = 6.42; auditory: M = 6.55) (F(1, 35) = 10.31, p < .01, η^2^p = .23), higher hit rates (visual: M = . 93; auditory: M = .88) (F(1, 35) = 9.45, p < .01, η^2^p = .21), and higher discrimination accuracy (visual: M = .93; auditory: M = .86) (F(1, 35) = 7.54, p < .01, η^2^p = .17). No interaction effects between group and task were observed. A repeated measures ANOVA (group × task) for RTs did not reveal significant differences between groups, suggesting that both groups took the same time responding to oddballs, while processing accuracy was higher in OAs (Figure 3).

**Figure 3.**
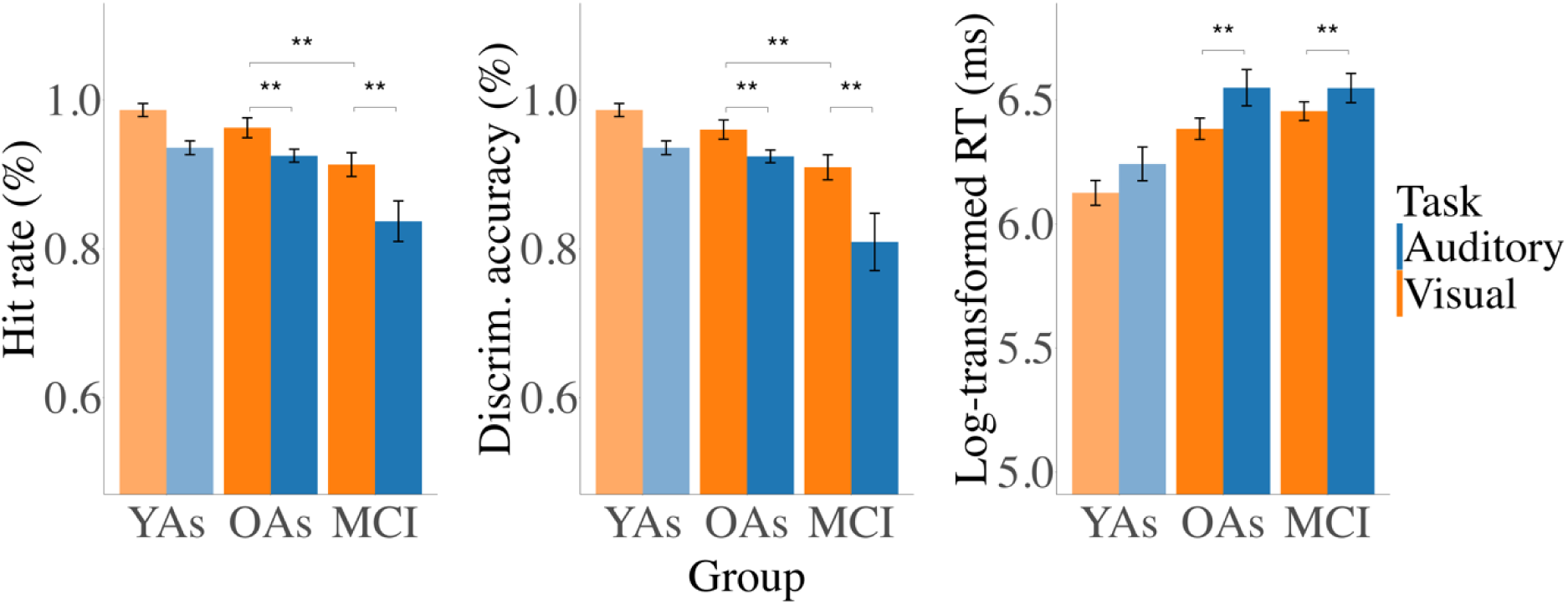
Mean percentage of hit rate, discrimination accuracy and RTs across two types of oddball tasks. Note that due to small sample sizes, data from YAs is not included in statistical analyses and included for illustrative purposes only (indicated by lighter shading). Error bars represent ±1 standard error. Significant differences are indicated by asterisks (**p < .01).

##### Within tasks

A t-test for hit rate revealed significantly better performance in OAs compared to MCI participants in the auditory oddball task, t(26.24) = 3.07, p < .01, d = .83, OAs (M = .92) and MCI (M = .83), as well as in the visual oddball task, t(35) = 2.14, p = .03, d = .72, OAs (M = .96) and MCI (M = .91). Similarly, a t-test for discrimination accuracy was significantly higher in OAs than in MCI in the auditory oddball task, t(24.12) = 2.91, p < .01, d = .77, OAs (M = .92) and MCI (M = .80) as well as in the visual oddball task, t(35) = 2.11, p = .04, d = .71, OAs (M = .96) and MCI (M = .90). A t-test on RTs for hits revealed no difference between groups in the auditory and visual oddball tasks.

#### 3.1.2 Pupillometry results

##### Between tasks

A repeated measures ANOVA (group x condition x task) for PD revealed the expected main effect of greater PD for oddball (M = 1.28) compared to standard stimuli (M = −.07) (F(1, 34) = 195.59, p < .001, η2p = .85) as well as a task x condition interaction effect (F(1, 34) = 27.24, p < .001, η2p = .44) (Figure 4), showing that PD was greater for oddball than standard stimuli in both visual (oddball: M = 1.07, standard: M = .07) (p < .001) and auditory task (oddball: M = 1.49, standard: M = −.22) (p < .001). For oddball stimuli, PD was higher in the auditory than the visual task (p < .01). For standard stimuli, higher PD was in the visual compared to auditory task (p < .01). Moreover, we found that the difference between PD for oddball and standard stimuli was greater on the auditory than the visual task (p < .001). No interaction between group and condition was found, indicating comparable pupil responses to different stimulus types for OAs and patients with MCI. No other interactions were observed. For the change in PD over time across the different conditions in the oddball task, see Figure 5.

**Figure 4.**
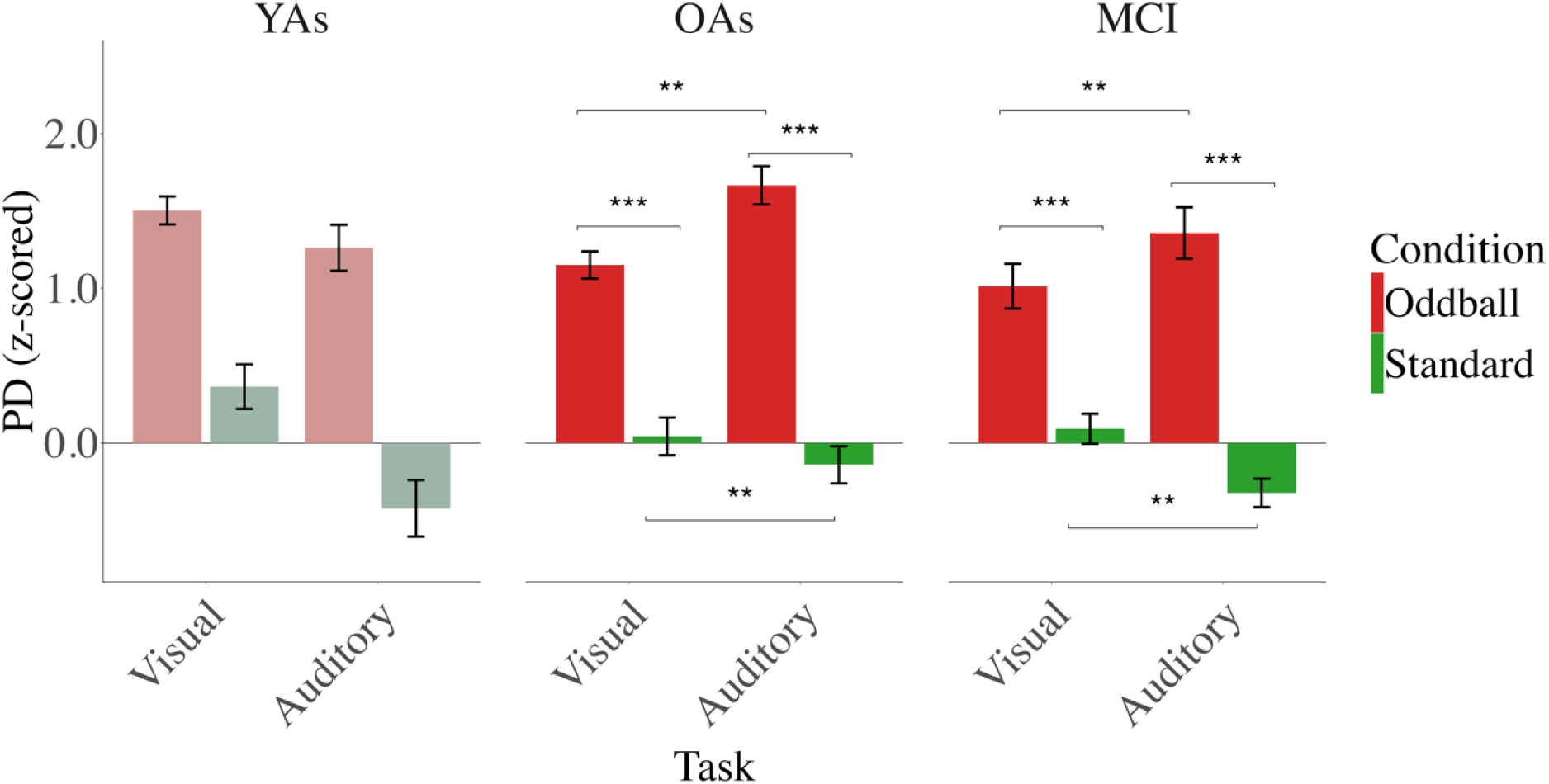
Mean PD across the visual and auditory oddball tasks, assessed as the mean in the time window (from 1 to 2 s and from .5 to 1.5 s after stimulus onset in the auditory oddball and visual oddball tasks, respectively). Note that due to small sample sizes, data from YAs is not included in statistical analyses and included for illustrative purposes only (indicated by lighter shading). Error bars represent ±1 standard error. Significant differences are indicated by asterisks (***p < .001, *p < .05).

**Figure 5.**
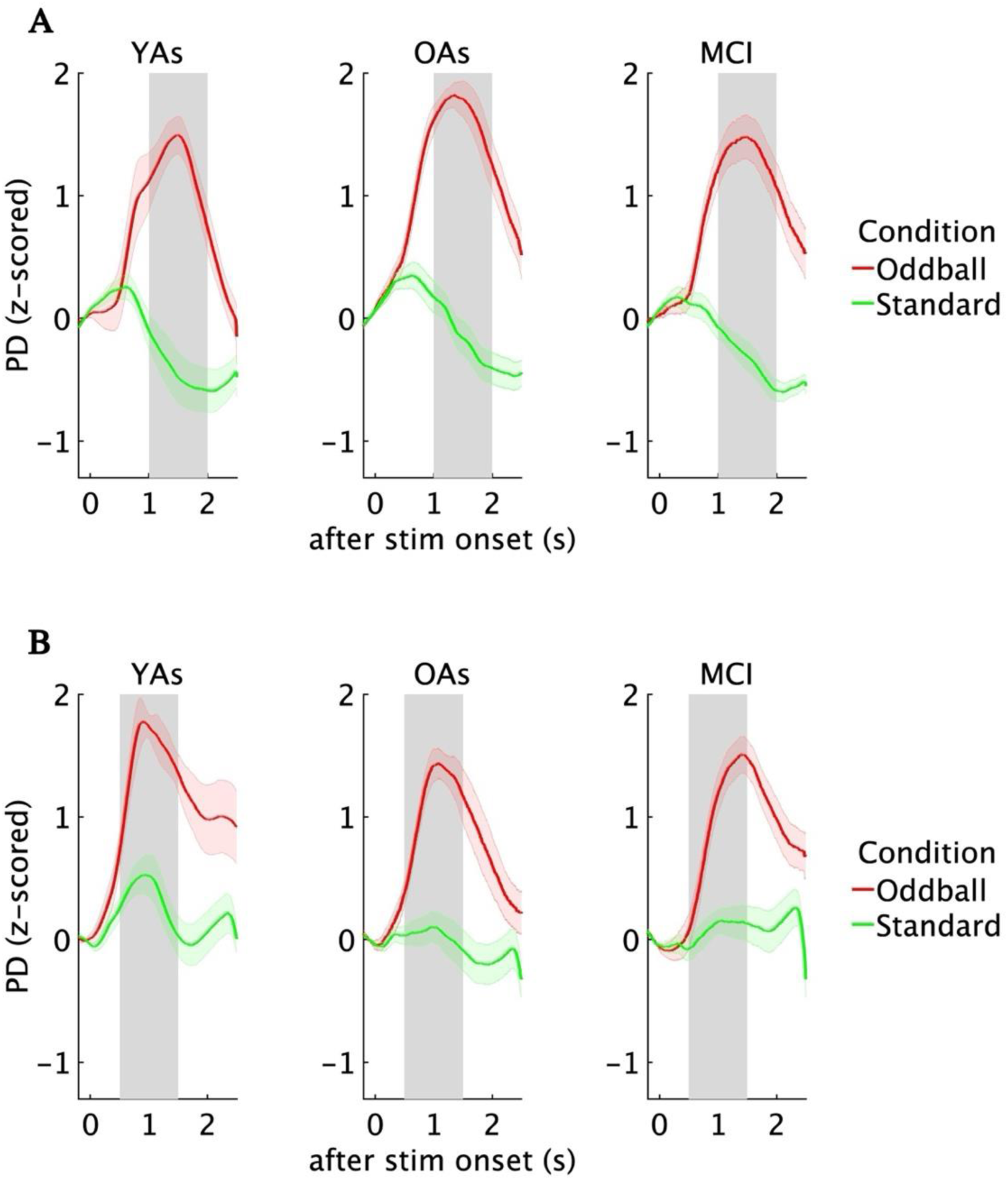
PD change over time in the oddball task. (A) PD change over time from 1 to 2 s after stimulus onset for oddball and standard **auditory oddball stimuli**. (B) PD change over time from .5 to 1.5 s after stimulus onset for oddball and standard **visual oddball stimuli**.

##### Within tasks

A repeated measures ANOVA (group x condition), analyzed separately for the visual and auditory oddball tasks, indicated a higher PD response for oddball stimuli (visual: M = 1.07; auditory: M = 1.49) compared to standard stimuli (visual: M = .07; auditory: M = −.22) in both the auditory oddball task (F(1, 34) = 176.10, p < .001, η2p = .83) and the visual oddball task (F(1, 34) = 85.41, p < .001, η2p = .71). Again, there were no main effect of the group and interactions between group and condition as well as between the baseline pupillary noise covariate and condition.

#### 3.1.3 Correlations between oddball task performance, pupil responses, cognitive screening measures as well as LC integrity

A significant correlation was observed between greater PD for the oddball stimuli and faster RTs for hits in both OAs, r = −.81, p < .01, and MCI patients, r = −.57, p < .01 on the visual task (Figure S6 in the the Supplement). This relationship was not apparent for standard stimuli in OAs, r = −.08, p = .78 or MCI patients, r = −.30, p = .18 (Figure S7 in the Supplement), suggesting that participants who can devote more attention and effort to the task-relevant stimuli may respond faster. A Williams’ t-test revealed that the correlation between PD and RTs was significantly stronger for PD for oddball stimuli (r = −.81) compared to PD for standard stimuli (r = −.08) in OAs, t(10) = −3.65, p < .01). However, although the correlation appeared to be stronger for oddball stimuli (r = −.57) compared to standard stimuli (r = −.30), in MCI patients, this difference did not reach statistical significance, t(18) = −1.08, p = .28. Because correlational coefficients for subsequent Williams’ t-tests should be calculated from the same sample, these correlational analyses were performed on data from the exact same individuals within each group (OAs: n = 13, MCI: n = 21). Furthermore, Fisher’s test showed that the association between PD for standard stimuli and RTs was not substantially stronger in OAs than in MCI patients, z = .57, p = .56. This pattern was not replicated in the auditory oddball task, as no significant correlations were observed between PD and behavioral data. In addition, attention scores from the ACE-R subtest did not correlate significantly with either PD or task performance in the oddball tasks. There were no significant correlations between LC integrity and PD in OAs and MCI patients.

### 3.2 Simon task

#### 3.2.1 Behavioral results

As YAs were fully included in assessments of the Simon task, statistical analyses include all three groups. A repeated measures ANOVA (group x condition) for accuracy revealed the expected main effect of higher accuracy for congruent trials (M = .96) as compared to incongruent trials (M = .93) (F(1, 59) = 10.42, p < .01, η^2^p = .15) and a group main effect (F(2, 59) = 5.52, p < .01, η^2^p = .16), with both YAs (M = .97) and OAs (M = .97) showing better accuracy than patients with MCI (M = .88; YAs vs MCI: p = .01; OAs vs MCI: p = .01), but no difference between YAs and OAs (p = 1.00). No significant group x condition interaction effect was observed (Figure 6A). A further one-way ANOVA for accuracy revealed a group effect for congruent trials (F(2, 59) = 6.53, p < .01, η^2^ = .18), showing both YAs (M = .99) and OAs (M = .99) had significantly better accuracy than MCI patients (M = .89; YAs vs MCI: p < .01; OAs vs MCI: < .01), with no difference between YAs and OAs (p = .99). For incongruent trials, a one-way ANOVA also showed a group effect (F(2, 59) = 3.63, p = .03, η^2^ = .11). However, pairwise comparisons showed no significant differences between groups (YAs vs OAs: p = .99, OAs vs MCI: p = .06), and the difference between YAs and MCI patients did not reach statistical significance (p = .052).

**Figure 6.**
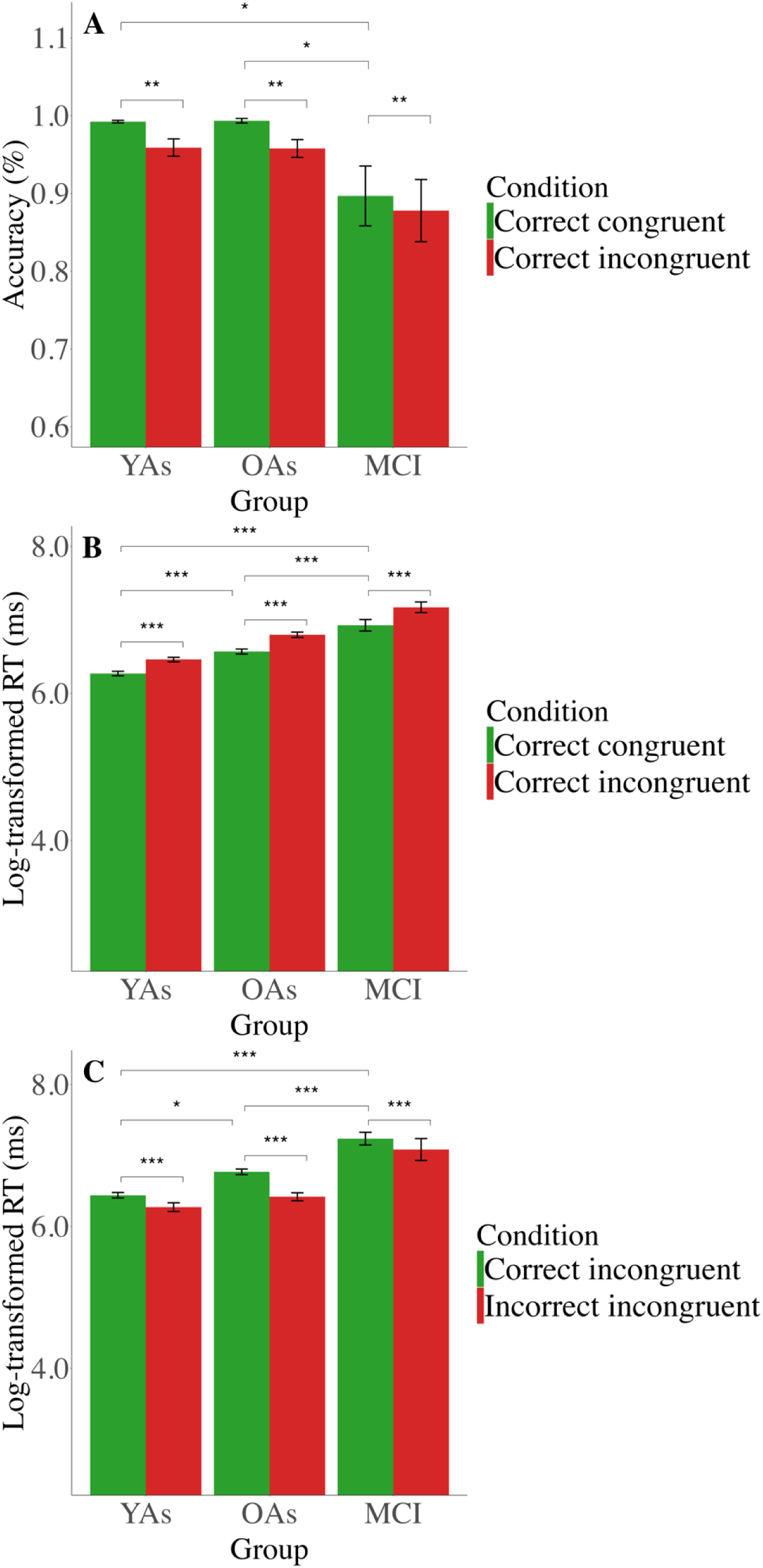
Behavioral performance in the Simon task. Error bars represent ±1 standard error. Significant differences are indicated by asterisks (*** p < .001, **p < .01, *p < .05). (A) Mean percentage of correct congruent and correct incongruent trials. (B) Mean RTs in correct congruent and correct incongruent trials. (C) Mean RTs in correct incongruent and incorrect incongruent trials.

A repeated measures ANOVA (group x condition) for RTs on correct congruent versus correct incongruent trials showed the expected main effect of condition (F(1, 59) = 313.04, p < .001, η^2^p = .84) and group (F(2, 59) = 50.00, p < .001, η^2^p = .63) (Figure 6B), with RTs being faster for correct congruent trials (M = 6.59) compared to correct incongruent trials (M = 6.81). The pattern of faster RTs during correct congruent trials is in line with enhanced cognitive control requirements on incongruent trials. Regarding group differences, MCI patients had slower RTs (M = 7.05) than both OAs (M = 6.68, p < .001) and YAs (M = 6.37, p < .001), and the latter two groups also showed significant differences between each other (p < .001). There was no significant group x stimulus type interaction effect for RTs on correct congruent versus correct incongruent trials. A one-way ANOVA for RTs found a group effect for congruent trials (F(2, 59) = 42.21, p < .001, η^2^ =.59), with MCI being the slowest (M = 6.93) compared to both YAs (M = 6.27, p < .001) and OAs (M = 6.57, p < .001), while OAs were slower on correct congruent trials compared to YAs (p < .001). For incongruent trials, a one-way ANOVA indicated a group effect (F(2, 59) = 53.75, p <.001, η^2^ =.64), with MCI being the slowest (M = 7.17) compared to both YAs (M = 6.46, p < .001) and OAs (M = 6.80, p < .001), and OAs were slower on correct incongruent trials compared to YAs (p < .001).

A repeated measures ANOVA (group x condition) for RTs on correct incongruent versus incorrect incongruent trials revealed significant main effects of condition (F(1, 37) = 28.15, p < .001, η^2^p = .43) and group (F(2, 37) = 33.52, p < .001, η^2^p = .64) (Figure 6C).

Specifically, incorrect incongruent responses (M = 6.59) were executed faster than correct incongruent responses (M = 6.82), consistent with the notion that incorrect responses often represent premature responses that are not inhibited by top-down motor control [63]. Patients with MCI had the slowest RTs (M = 7.16) compared to both YAs (p < .001) and OAs (p < .001), while OAs (M = 6.60) were slower than YAs (M = 6.36, p = .04). There was no significant group x stimulus type interaction effect for RTs on correct incongruent versus incorrect incongruent trials.

#### 3.2.2 Pupillometry results

The repeated measures ANOVA (group x condition) for PD showed the expected main effect of condition (F(1, 58) = 73.47, p < .001, η2p = .55), with higher PD for incongruent (M = .62) versus congruent trials (M = .002), suggesting more attention and/ or effort when processing incongruent stimuli. A condition x group interaction (F(2, 58) = 3.22, p = .04, η2p = .10) indicated greater PD for incongruent (M = .65) versus congruent trials (M = −.09) in OAs (p < .001) and for incongruent (M = .55) versus congruent (M = −.25) trials in MCI patients (p < .001), whereas YAs somewhat unexpectedly did not show a significant difference between congruent (M = .34) and incongruent conditions (M = .67) (Figure 7). Additionally, PD was found to be greater for congruent trials in YAs than in MCI patients (p = .03). No significant interaction was found between the baseline pupillary noise covariate and condition. For the change in PD over time across the different conditions in the Simon task, see Figure 8.

**Figure 7.**
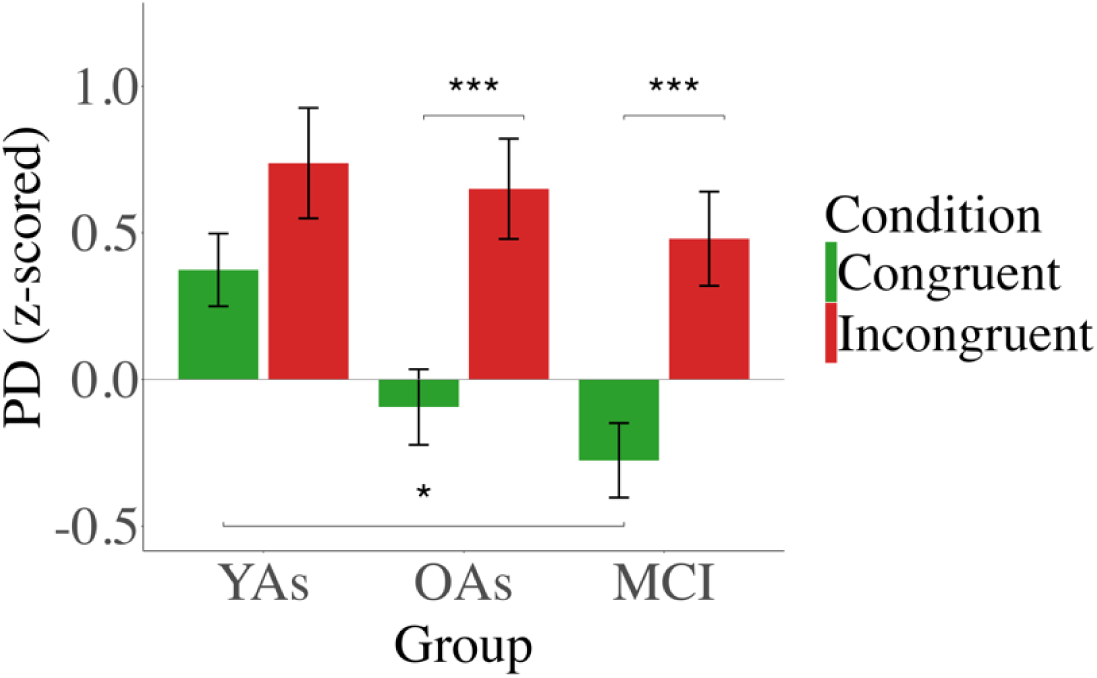
Mean PD in the Simon task, assessed as the mean in the time window (from 1 to 2 s after stimulus onset). Error bars represent ±1 standard error. Significant differences are indicated by asterisks (***p < .001, *p < .05).

**Figure 8.**
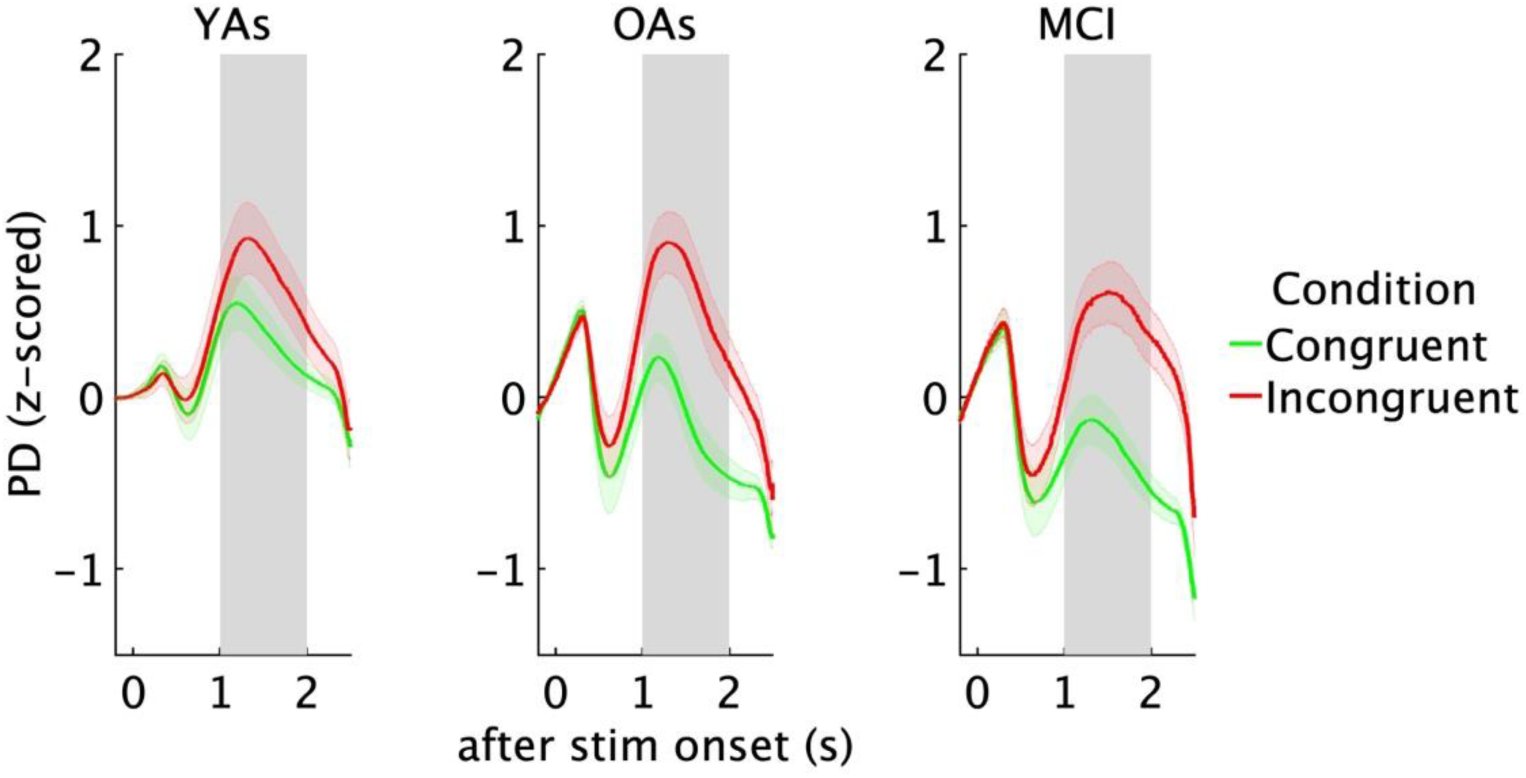
PD change over time from 1 to 2 s after stimulus onset for congruent and incongruent Simon task stimuli.

#### 3.2.3 Correlations between oddball task performance, pupil responses, cognitive screening measures as well as LC integrity

Examining the correlations between behavioral data and PD revealed faster RTs for correctly performed incongruent trials in YAs which showed a greater PD on incongruent trials, r = −.63, p < .01 (Figure S8 in the Supplement), suggesting that YAs who expended more cognitive control responded faster. Second, faster RTs on incorrectly performed incongruent trials were associated with increased PD on incongruent trials in YAs, r = −.61, p = .02 (Figure S9 in the Supplement). Increased PD on erroneous incongruent trials might reflect increased (albeit unsuccessful) efforts on incongruent trials, or heightened attentional arousal in the context of a typically conscious action slip error [63]. The same was observed for MCI patients, r = −.82, p < .01 (Figure S9 in the Supplement). In addition, we found that a more pronounced Simon effect (i.e., PD incongruent minus PD congruent) correlated with a faster RTs on correctly performed incongruent trials in YAs, r = −.62, p < .01 (Figure S10 in the Supplement).

To determine whether interindividual differences in attention and memory capacity contributed to interindividual differences in the Simon task, we correlated attention and memory scores from the ACE-R subtests with PD and behavioral data from the Simon task. We found that individuals with higher memory capacity showed faster RTs for correct congruent trials in MCI patients, r = −.52, p = .02 (Figure S11 in the Supplement). Similarly, higher memory scores were correlated with better performance on correct congruent, r = .64, p < .01 (Figure S12 in the Supplement) and incongruent trials in the MCI group, r = .46, p = .03 (Figure S13 in the Supplement). Similar results were observed for the MMSE: higher memory scores were associated with a higher percentage of correctly completed congruent trials in the MCI group, r = .51, p = .02 (Figure S14 in the Supplement). Linear regression analyses did not reveal significant associations between LC integrity and PD in the Simon task in OAs and MCI patients. However, we found that in MCI patients, higher LC integrity was associated with faster RTs for correct incongruent trials (b = −5.26, p = .01) (Figure S15 in the Supplement).

## 4. DISCUSSION

The results of the study showed that the expected involvement of PD in top-down attentional modulation or cognitive effort during the oddball and Simon tasks occurs to a similar extent in both elderly groups, i.e., healthy OAs and MCI patients. For the oddball task, the oddball effect, represented by increased PD for oddball stimuli, was replicated in OAs and MCI patients. Performance was better in OAs than in MCI patients, as indicated by a higher hit rate for oddball stimuli and discrimination accuracy in OAs compared to MCI patients. For the Simon task, PD was increased for incongruent as compared to congruent trials in OAs and MCI patients, but not in YAs. A classic Simon effect was observed, with incongruent trials eliciting lower accuracy and longer RTs, and MCI patients performing more slowly and less accurately than OAs. Higher PD responses were associated with faster RTs in all tasks. Simon’s task performance in MCI patients was linked to memory scores on the ACE-R and MMSE.

The behavioral results of the oddball task revealed a mixed pattern of attentional resource allocation in MCI. MCI patients showed intact motor response speed with comparable RTs to healthy OAs in both oddball tasks. However, their ability to sustain attention and suppress irrelevant task stimuli is impaired, with lower accuracy levels, consistent with studies suggesting that executive function deficits are a significant predictor of the transition from MCI to AD [64]. Importantly, OAs and MCI patients showed greater PD to oddball stimuli compared to standard stimuli. This PD-based oddball effect is well-established in auditory and visual modalities [65,8]. Although we expected a reduced attentional allocation in MCI versus healthy aging, represented by larger PD in OAs, we did not observe any group differences in different conditions. Furthermore, although the results must be treated with caution given the limited sample size (Figures S1-S3 in the Supplement) in both groups, individuals with greater PD to oddball stimuli reacted faster to oddball stimuli, supporting that PD reflects attentional allocation and motor effort [2,44]. Conversely, a lack of correlation with standard stimuli may suggest reduced processing resource needs. Thus, while MCI patients show expected behavioral decline in attentional modulation, PD can serve as an informative marker of executive control of attention and motor effort in both healthy OAs and MCI patients.

An unexpected finding was a larger PD response to oddball stimuli in the auditory than in the visual task. Since PD should reflect cognitive effort, this finding may be related to the fact that the auditory oddball task is more challenging than the visual oddball task. Indeed, both OAs and MCI patients performed more accurately and faster on visual than auditory oddball stimuli. A possible explanation is that age-related visual acuity decline was better regulated than potential hearing impairments in the auditory task. All participants completed a visual acuity test and received visual aids if needed. Participants with hearing impairments could request louder volume, but no standard hearing tests were performed. As a result, interindividual differences in the ease of processing the stimuli may have contributed to the variance in executive processes reflected in PD. This additional variance may have masked the interindividual relationship between greater PD and faster RTs for oddball stimuli that was only observed in the visual task. Both the oddball tasks proved useful in identifying behavioral and PD-related executive processes in MCI, but future studies should control for the sensory accessibility of stimuli to improve the cognitive characterization of MCI by PD.

While both oddball stimuli in the oddball task and incongruent stimuli in the Simon task can be argued to reflect higher attentional demands and motor execution effort, the need to overcome incongruent response tendencies in the Simon task [34] presents an additional demand for executive control. In this sense, comparing these task types in the same population allowed us to assess whether both attentional and executive control can be examined using pupillometry in MCI. Our study replicated the Simon effect [66], showing reduced accuracy and slower RTs during incongruent trials. Furthermore, during incorrect incongruent trials, RTs were faster than during correct ones, supporting the idea that conflict errors represent ‘action slips’ [63]. While there was no interaction effect, which suggests a similar effect of condition on behavioral performance across all groups, MCI patients showed slower RTs specifically on incongruent trials than YAs and OAs, despite maintaining comparable accuracy. This pattern of results suggests that during incongruent trials, MCI patients, by taking additional time, allocate additional cognitive resources to maintain accuracy. They are slower and less accurate during easier congruent trials, indicating inconsistent attentional engagement across both conditions. Although our analyses revealed similar accuracy levels for YAs and OAs during both trial types, OAs exhibited slower RTs compared to YAs. This indicates the decrease in processing speed with aging [67], while resource allocation remains intact.

As expected, PD was increased for incongruent trials than for congruent trials in all groups, suggesting greater top-down attentional control and cognitive effort invested in the incongruent condition [44]. OAs and MCI patients showed enhanced PD responses for incongruent trials, while YAs showed no difference in PD between trial types. This is unexpected, as YAs are assumed to allocate effort and attention selectively in response to increased task demands [7]. Inspecting individual response patterns (Figure S4 in the Supplement) revealed that most YAs (15 out of 22) showed the expected larger PD for incongruent than congruent trials. Studies with larger sample sizes are needed to assess whether interindividual differences in incongruency-specific PD effects in YAs can identify different motivational levels or task strategies.

While MCI patients showed worse accuracy and RTs than OAs across congruent and incongruent conditions, PD responses in the Simon task were similar between groups. As in the oddball task, examining links of interindividual differences in PD and task performance needs to be interpreted cautiously, given the small sample sizes (Figures S4 and S5 in the Supplement). Nonetheless, some consistent patterns emerged with individuals with faster RTs generally showing larger PDs, both on correct and incorrect incongruent trials (Figures S8 and S9 in Supplement) as well as about the differences in PD correct incongruent – correct congruent (Figure S10 in the Supplement) reflecting a more specific measure of increased PD due to increased attentional and cognitive control. This was rarely the case for OAs, which might suggest less heterogeneity in this group in mobilizing additional cognitive resources. Finally, MCI patients were naturally the only group with sufficient variance in cognitive dementia screenings for examining interindividual differences. Higher accuracy and lower RTs in the Simon task were linked to better memory scores, underlining the functional relevance of examining executive function in MCI.

Unexpectedly, we observed increased dilation and constriction in the early pupillary processes after stimulus presentation in OAs and MCI in the Simon task. It is presently unclear what these early pupil reactions reflect and why they differ across age groups. If earlier time windows in pupil response reflect earlier attentional and sensory processes, as appears to be the case for EEG data [68], this might indicate heightened sensory processing in older groups. Future studies should explore these early pupil reactions in varying sensory processing demands.

Finally, on neither task, changes in PD measured were associated with LC integrity, even though LC integrity was lower in MCI compared to YAs and OAs. The LC, a key norepinephrine hub, is involved in attentional regulation processes [3,5], and LC firing has been linked to increased PD [22,13,14]. However, PD response to salient stimuli may be influenced by multiple neural pathways and thus not exclusively related to the noradrenergic LC system [69]. Another possible explanation is that compensatory efforts may add degrees of freedom in functional correlates, making it more challenging to detect direct relationships with LC integrity. To address whether and under which conditions PD can serve as an indicator for LC function in aging and MCI/ AD, larger data acquisitions of reliable assessments in LC fMRI and concurrent PD are needed. We observed a negative relationship between LC integrity and RTs during correct incongruent trials in the Simon task in MCI patients, suggestive of higher executive control in individuals with more intact LCs. Previous research links LC integrity and RTs in executive tasks [70] and other cognitive paradigms [35,36,37], highlighting the relevance of LC decline in neurodegeneration.

Taken together, our study suggests that PD measures of the Simon and oddball tasks can assess cognitive function changes related to executive and attentional processing in healthy aging and MCI. Consistent with our hypothesis, OAs and MCI patients showed a similar pattern of pupillometric results, with oddball stimuli and incongruent trials requiring more attention/effort than standard stimuli in the oddball task and congruent trials in the Simon task, respectively. The preservation of similar PD response patterns across conditions, coupled with similar correlation patterns across YAs, OAs, and MCI patients, suggests that basic neural mechanisms governing attentional resource allocation remain intact despite behavioral performance differences and can be exploited using PD in early neurodegenerative stages.

## ACKNOWLEDGMENTS

We would like to thank the participants, caregivers, and researchers who were part of this study.

## CONSENT STATEMENT

Written informed consent was obtained from all participants and their carers prior to participation in the experiment.

