## Supplemental Table S1 for "Pupil dilation as a marker of attention/effort in aging and mild cognitive impairment"

*Table S1*. Descriptive statistics of the study participants

| **Group** | **YAs** | **OAs** | **MCI** | **p-value** |
| --- | --- | --- | --- | --- |
| N | 30 | 30 | 33 |  |
| Gender (M/ F) | 12/18 | 9/21 | 22/11 | .01 |
| Mean age (SD) | 23.07 (2.75) ^*, †^ | 71.37 (6.34) ^*^ | 74.21 (7.28) ^†^ | < .001 |
| Age range | 20-30 | 60-82 | 60-85 |  |
| Native English speakers | 60% ^*, †^ | 97% ^*^ | 100% ^†^ | < .001 |
| Mean ACE-R total | 94.23 (4.75) ^†^ | 96.03 (3.37) ^‡^ | 78.58 (10.96) ^†, ‡^ | < .001 |
| Mean ACE-R memory (SD) | 24.27 (2.11) ^†^ | 24.63 (2.00) ^‡^ | 15.88 (6.04) ^†, ‡^ | < .001 |
| Mean ACE-R attention/ orientation (SD) | 17.87 (.43) ^†^ | 17.80 (.48) ^‡^ | 15.24 (1.90) ^†, ‡^ | < .001 |
| Mean ACE-R verbal fluency | 12.57 (1.43) ^†^ | 12.93 (1.14) ^‡^ | 9.43 (2.63) ^†, ‡^ | < .001 |
| Mean ACE-R language | 24.17 (2.15) ^*^ | 25.53 (.77) ^*, ‡^ | 24.07 (2.23) ^‡^ | < .01 |
| Mean ACE-R visuospatial ability | 15.43 (1.13) ^†^ | 15.20 (1.09) ^‡^ | 13.98 (2.56) ^†, ‡^ | < .01 |
| Mean MMSE (SD) | 29.53 (1.07) ^†^ | 29.43 (.72) ^‡^ | 25.18 (2.70) ^†, ‡^ | < .001 |
| LC integrity (SD) | .13 (.03) ^*, †^ | .16 (.04) ^*, ‡^ | .10 (.03) ^†, ‡^ | < .001 |
| *Note.* Descriptive statistics include number of participants (N), distribution of gender by group, mean age and age range in years, percentage of native speakers, mean ACE-R total score with a maximum of 100, mean ACE-R score in the memory domain with a maximum of 26, mean ACE-R score in the attention/orientation domain with a maximum of 18, mean ACE-R score in the verbal fluency with a maximum of 14, mean ACE-R score in the language with a maximum of 26, mean ACE-R score in the visuospatial ability with a maximum of 16, mean MMSE score on a scale of 0 (poor) to 30 (good), and mean LC integrity. SD – standard deviation. P-values for group comparisons based on one-way ANOVAs for comparisons of three participant groups and independent samples t-test for comparisons of two participant groups. To evaluate group differences for categorical variables, chi-squared tests of independence (χ²) was used. Significance levels were set at α = .05. ^*^ – indicates significant differences between younger adults (YAs) and older adults (OAs). ^†^ – indicates significant differences between YAs and patients with mild cognitive impairment (MCI). ^‡^ – indicates significant differences between OAs and MCI patients. | | | | |
