## Supplemental Table S2 for "Pupil dilation as a marker of attention/effort in aging and mild cognitive impairment"

Table S2. Mean number of included PD trials by task, condition, and group

| **Task/ Group** | **YAs** | **OAs** | **MCI** | **p-value** |
| --- | --- | --- | --- | --- |
| **Simon task** | (n = 22) | (n = 20) | (n = 20) |  |
| Congruent trials | 127 | 133 | 111 | .07 |
| Incongruent | 32 | 33 | 29 | .18 |
| **Auditory oddball task** | (n = 7) | (n = 14) | (n = 23) |  |
| Oddball | 13 | 12 | 10 | .08 |
| Standard | 48 | 36 | 37 | .13 |
| **Visual oddball task** | (n = 7) | (n = 14) | (n = 23) |  |
| Oddball | 15^*^ | 11^*^ | 12 | .03 |
| Standard | 48 | 37 | 39 | .17 |
| Note. Values represent the mean number of trials included per condition. P-values for group comparisons based on one-way ANOVAs for comparisons of three participant groups. Significance levels were set at α = .05. ^*^ – indicates significant differences between younger adults (YAs) and older adults (OAs). There were no significant differences between any groups and patients with mild cognitive impairment (MCI). | | | | |
