## Supplemental Figure S1 for "Pupil dilation as a marker of attention/effort in aging and mild cognitive impairment"

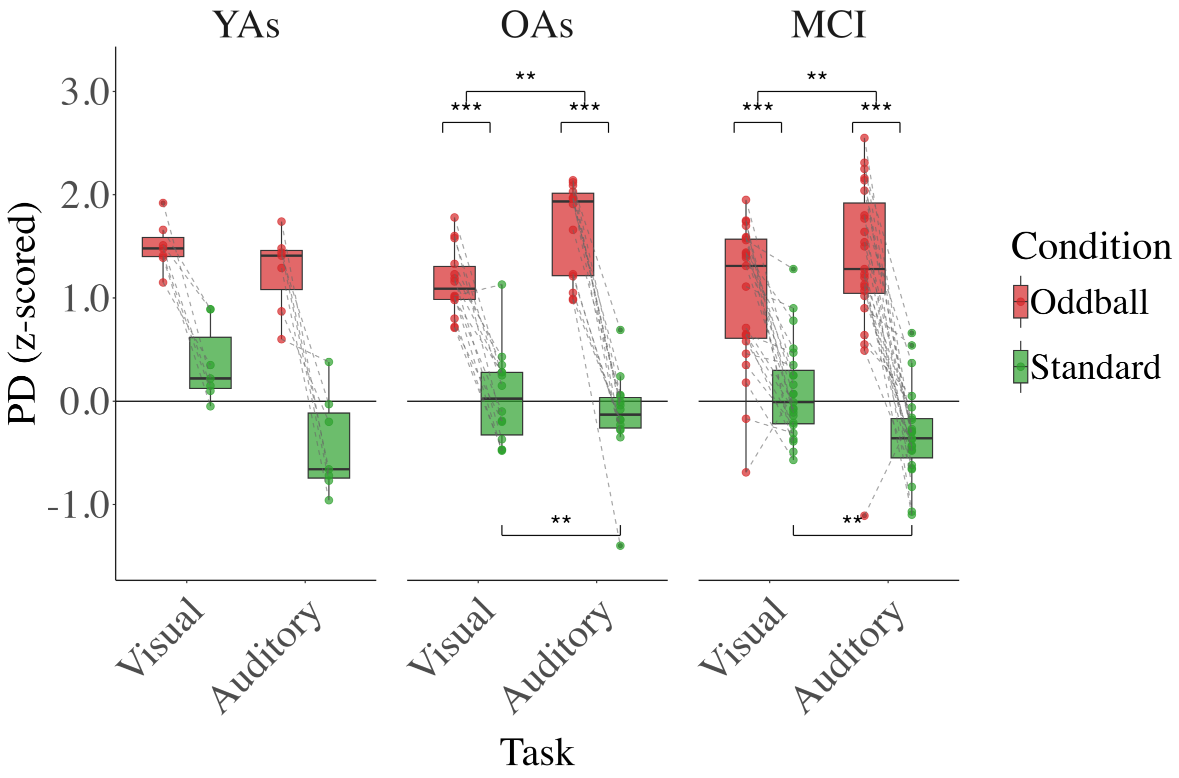


*Figure S1*. Overview of pupil dilation (PD) results in the oddball tasks for individual participants (younger adults (YAs) (n=7), older adults (OAs) (n=14), and patients with mild cognitive impairment (MCI) (n=23)). Data are presented as standardized z-scores, with box plots showing the median (horizontal line), interquartile range (box), and whiskers extending 1.5 × interquartile range from the quartiles. Each point represents an individual participant's mean PD, with dashed gray lines connecting paired observations from the same individual across conditions. Significant differences are indicated by asterisks (***p < .001, *p < .05). The pattern of directional differences in PD between oddball and standard stimuli was examined across two participant groups (OAs and MCI patients), with a meaningful difference defined as a relative difference greater than 10%, calculated as the absolute difference between conditions divided by the absolute value of the mean of the two conditions. Chi-square goodness-of-fit tests indicated that the observed distribution was significantly different from a uniform distribution in all groups, suggesting that oddball trials typically elicit greater PD than standard stimuli across both oddball tasks. In the visual task, YAs: 7 participants showed greater PD for oddball stimuli, 0 showed greater PD for standard stimuli, and 0 showed no meaningful difference ($\chi^2(2) = 14$, $p < .001$). OAs: 13 participants showed greater PD for oddball stimuli, 1 showed greater PD for standard stimuli, and 0 showed no meaningful difference ($\chi^2(2) = 22.42$, $p < .001$). MCI patients: 22 participants showed greater PD for oddball stimuli, 1 showed greater PD for standard stimuli, and 0 showed no meaningful difference ($\chi^2(2) = 40.26$, $p < .001$). In the auditory task, YAs: 7 participants showed greater PD for oddball stimuli, 0 showed greater PD for standard stimuli, and 0 showed no meaningful difference ($\chi^2(2) = 14$, $p < .001$). OAs: 14 participants showed greater PD for oddball stimuli, 0 showed greater PD for standard stimuli, and 0 showed no meaningful difference ($\chi^2(2) = 28$, $p < .001$). MCI patients: 22 participants showed greater PD for oddball stimuli, 1 showed greater PD for standard stimuli, and 0 showed no meaningful difference ($\chi^2(2) = 40.26$, $p < .001$).
