## Supplemental Figure S2 for "Pupil dilation as a marker of attention/effort in aging and mild cognitive impairment"

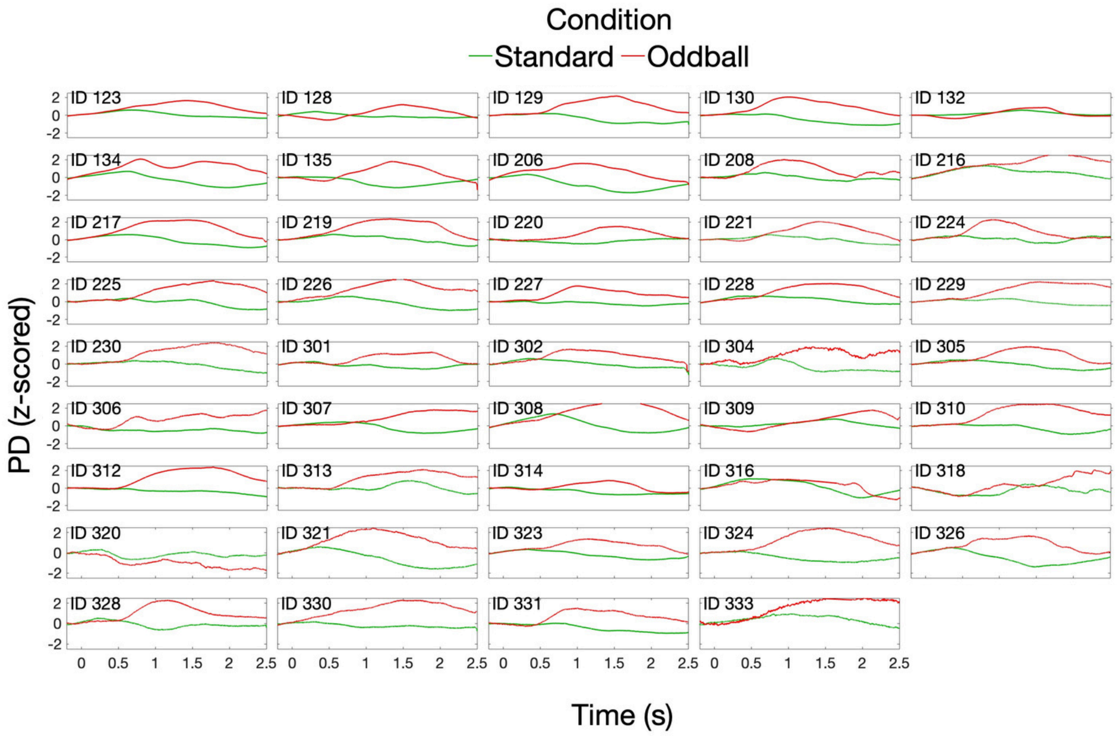


*Figure S2*. Averaged pupil dilation (PD) during trials performed in standard (green) and oddball (red) conditions per individual in the **auditory oddball task**. Younger adults have ID numbers ranging from 100 to 199, older adults have ID numbers ranging from 200 to 299, and patients with mild cognitive impairment have ID numbers starting from 300.
