## Supplemental Figure S4 for "Pupil dilation as a marker of attention/effort in aging and mild cognitive impairment"

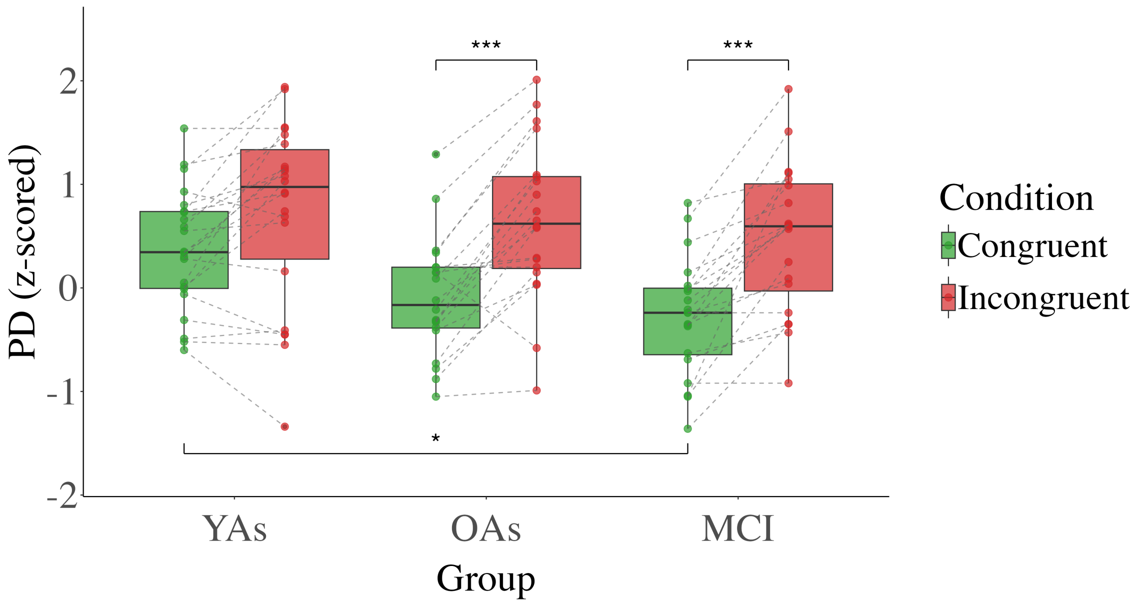


*Figure S4*. Overview of pupil dilation (PD) results in the Simon task for individual participants (younger adults (YAs) (n=22), older adults (OAs) (n=20), and patients with mild cognitive impairment (MCI) (n=20)). Data are presented as standardized z-scores, with box plots showing the median (horizontal line), interquartile range (box), and whiskers extending 1.5 × interquartile range from the quartiles. Each point represents an individual participant's mean PD, with dashed gray lines connecting paired observations from the same individual across conditions. Significant differences are indicated by asterisks (***p <. 001). The pattern of directional differences in PD between congruent and incongruent trials was examined across all three participant groups, with a meaningful difference defined as a relative difference greater than 10%, calculated as the absolute difference between conditions divided by the absolute value of the mean of the two conditions. Chi-square goodness-of-fit tests indicated that the observed distribution was significantly different from a uniform distribution in all groups, suggesting that incongruent trials typically elicit greater PD than congruent trials. YAs: 15 participants showed greater PD for incongruent trials, 5 showed greater PD for congruent trials, and 2 showed no meaningful difference ($\chi^2(2) = 12.63$, $p < .01$). OAs: 17 participants showed greater PD for incongruent trials, 1 showed greater PD for congruent trials and 2 showed no meaningful difference ($\chi^2(2) = 24.1$, $p < .001$). MCI patients: 18 participants showed greater PD for incongruent trials, 0 greater PD for congruent trials, and 2 showed no meaningful difference ($\chi^2(2) = 29.2$, $p < .001$).
