## Supplemental Figure S7 for "Pupil dilation as a marker of attention/effort in aging and mild cognitive impairment"

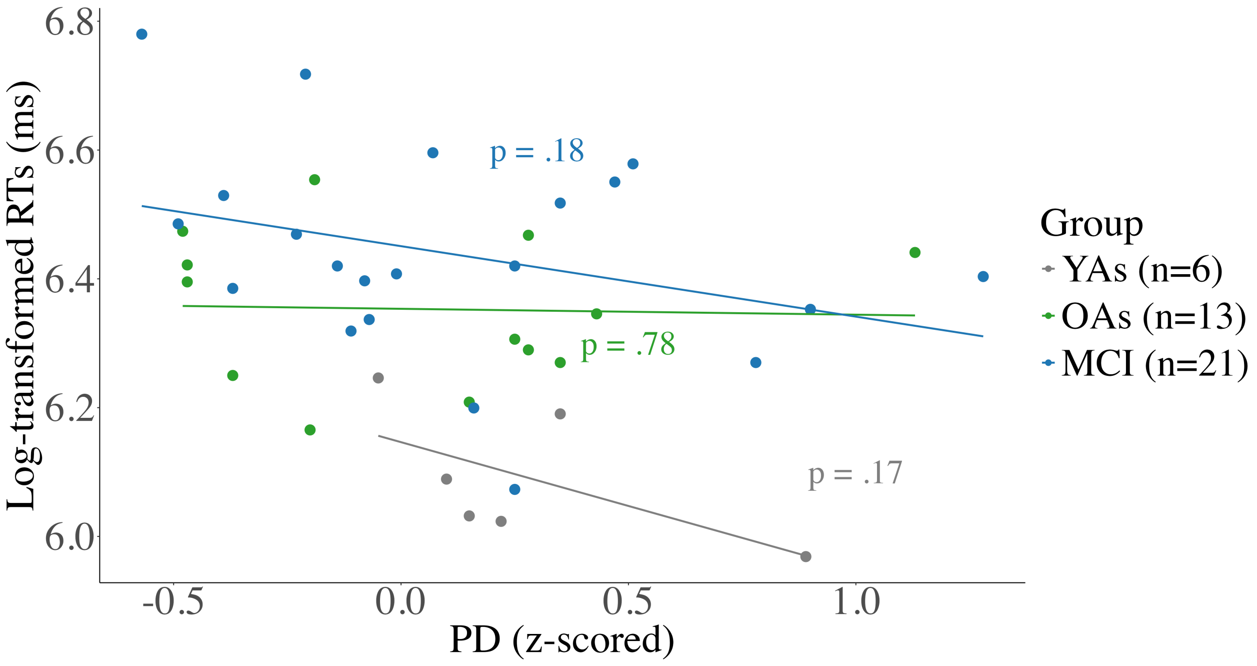


*Figure S7*. Spearman correlation between pupil dilation (PD) for the **standard stimuli** and hit reaction times (RTs) across groups in the **visual oddball task.** YAs – younger adults, OAs – older adults, MCI – patients with mild cognitive impairment.
