## Supplemental Figure S9 for "Pupil dilation as a marker of attention/effort in aging and mild cognitive impairment"

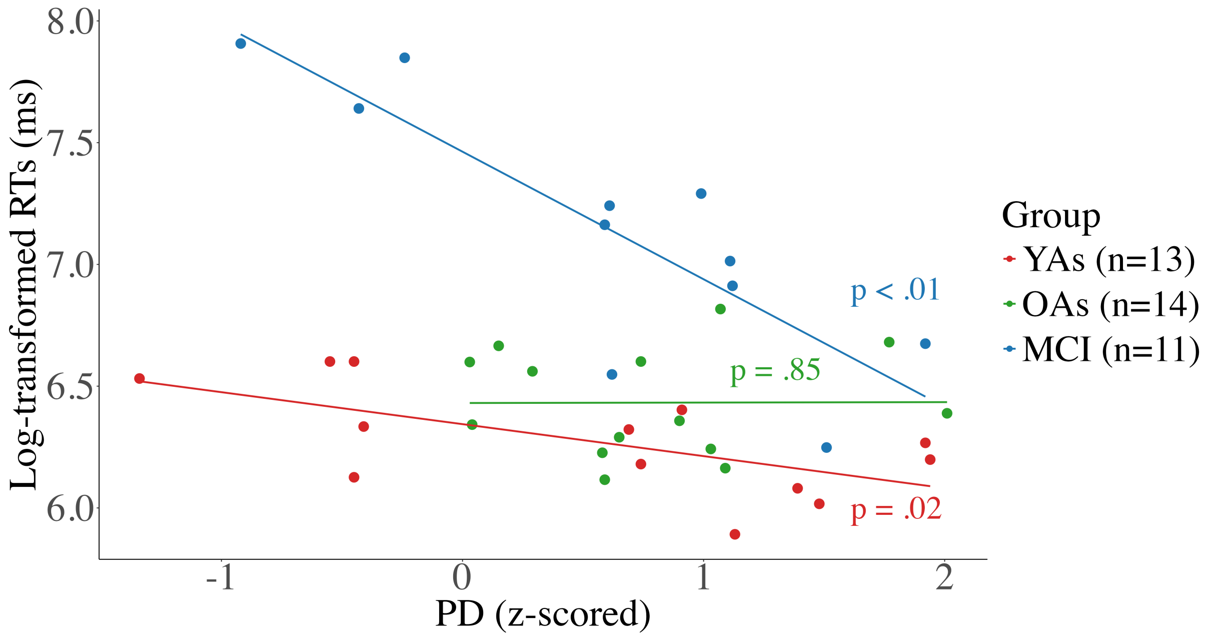


*Figure S9*. Spearman correlation between pupil dilation (PD) and reaction times (RTs) in incorrect incongruent trials across groups in the Simon task. YAs – younger adults, OAs – older adults, MCI – patients with mild cognitive impairment.
