## Supplemental Figure S10 for "Pupil dilation as a marker of attention/effort in aging and mild cognitive impairment"

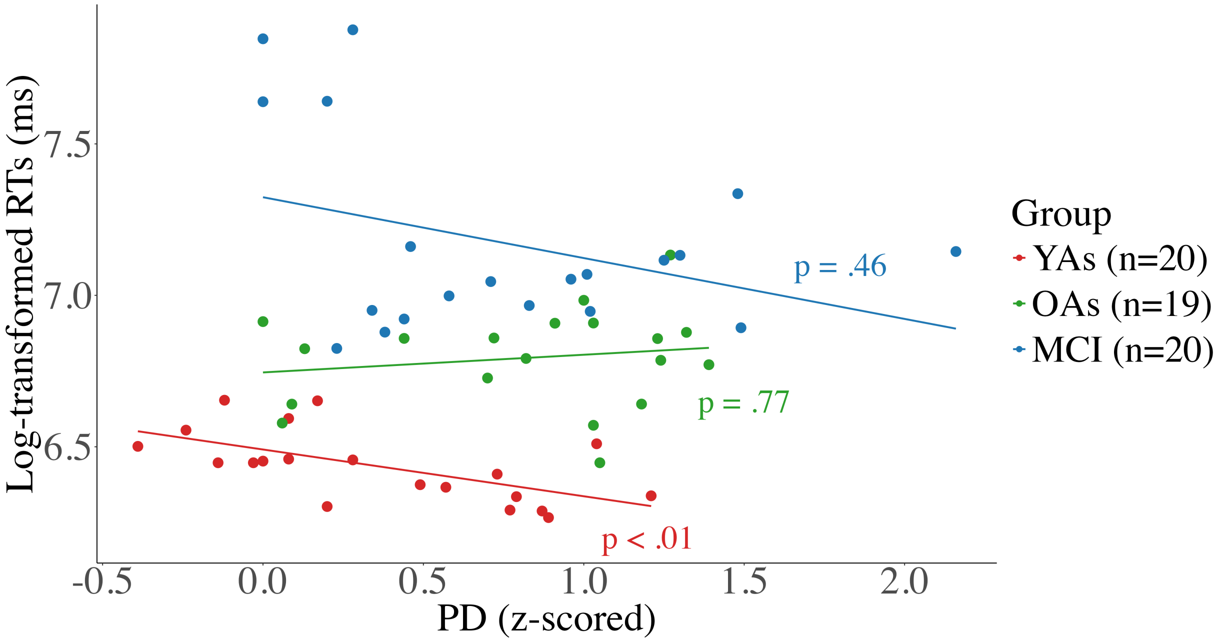


*Figure S10*. Spearman correlation between pupil dilation (PD) difference (PD in correct incongruent trials minus PD in correct congruent trials) and reaction times (RTs) in correct incongruent trials across groups in the Simon task. YAs – younger adults, OAs – older adults, MCI – patients with mild cognitive impairment.
