## Supplemental Figure S12 for "Pupil dilation as a marker of attention/effort in aging and mild cognitive impairment"

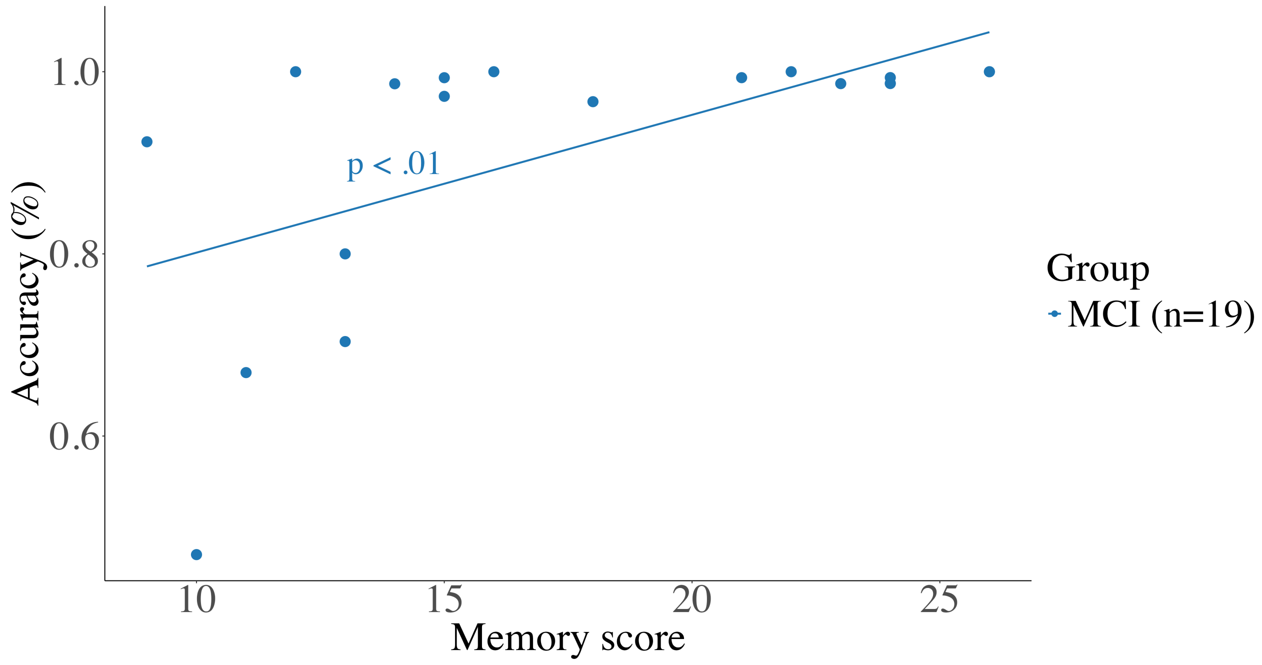


*Figure S12*. Spearman correlation between memory score in the revised version of Addenbrooke's Cognitive Examination and accuracy in correct congruent trials in the Simon task. MCI – patients with mild cognitive impairment.
