## Supplemental Figure S14 for "Pupil dilation as a marker of attention/effort in aging and mild cognitive impairment"

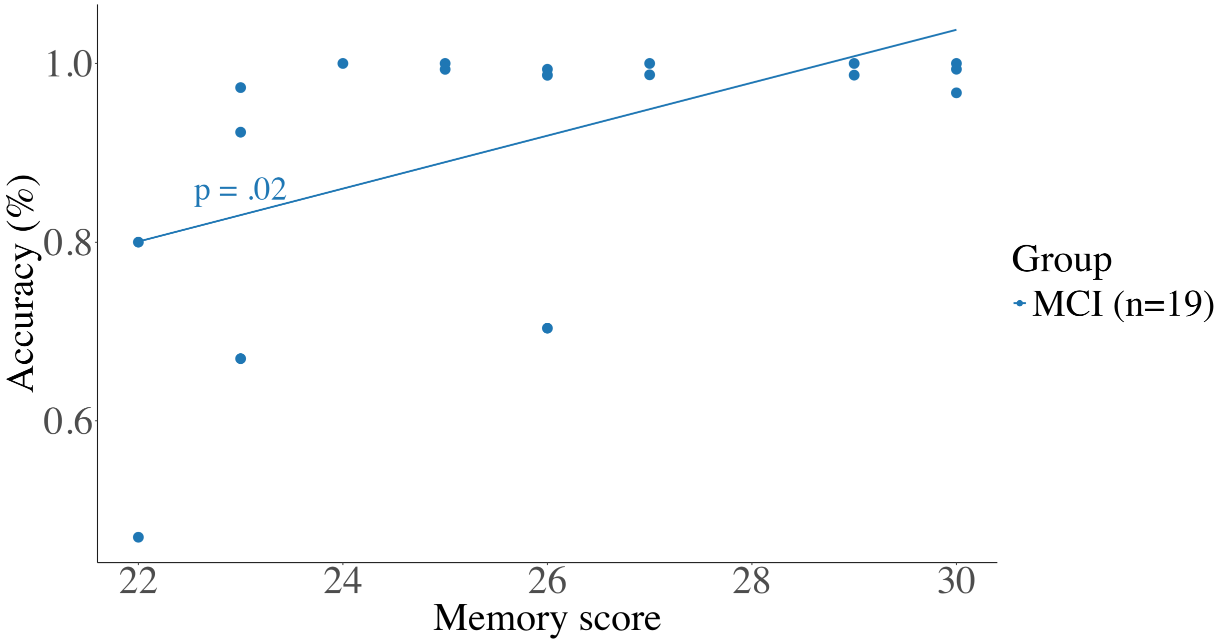


*Figure S14*. Spearman correlation between memory score in the Mini Mental State Examination and accuracy in correct congruent trials in the Simon task. MCI – patients with mild cognitive impairment.
