## Supplemental Figure S15 for "Pupil dilation as a marker of attention/effort in aging and mild cognitive impairment"

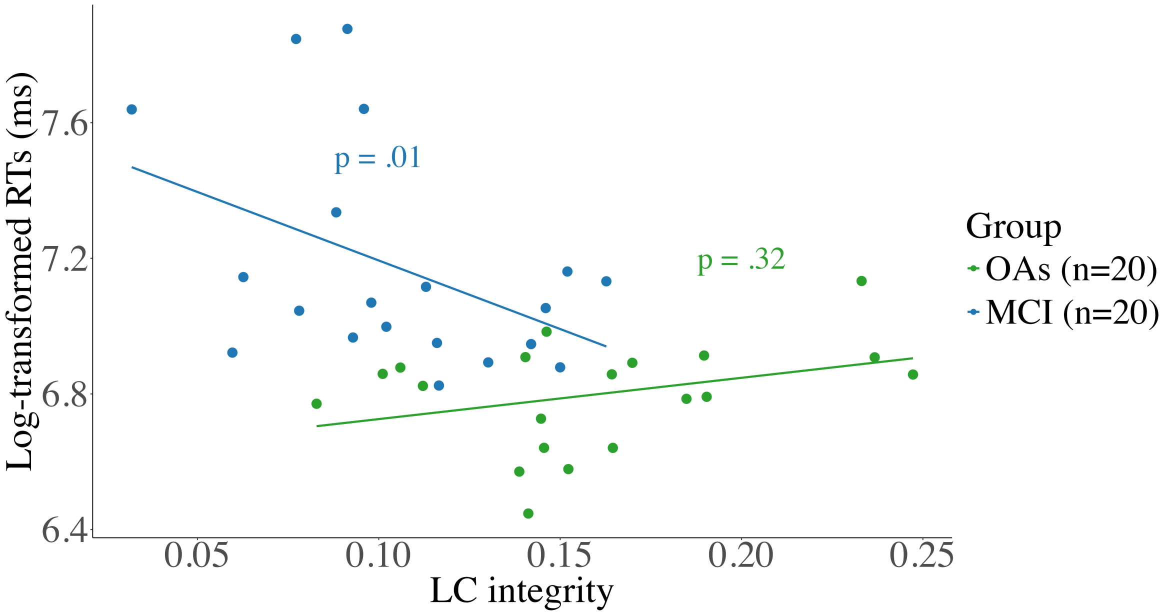


*Figure S15*. Relationship between locus coeruleus (LC) integrity and reaction times (RTs) in correct incongruent trials across groups in the Simon task. OAs – older adults, MCI – patients with mild cognitive impairment.
